# ExTaSy: A swappable CRISPR platform for endogenous tagging in *Drosophila melanogaster*

**DOI:** 10.64898/2026.01.16.699669

**Authors:** Sean Hubbert, Zara Valliji, Sadeen Abu-Hananah, Sarah Phal, Marina Luchner, Emma Watts, Giulia Biasi, Korneel Hens, Sebastian Kittelmann

## Abstract

Gene tagging enables functional analysis of proteins *in vivo*. Currently existing technologies in *Drosophila* suffer from drawbacks including limited flexibility, copy number variation, and/or imprecise expression. To address this, we have developed the Exchangeable Tagging System (ExTaSy), a CRISPR/Cas9-based platform which introduces a 3XHA tag into the endogenous gene locus. Importantly, the 3XHA can be subsequently exchanged for other tags using fly crosses, making the platform highly versatile and accessible. Simultaneously, an excisable transgenic marker allows virtually scarless locus modification. Here, we report successful tagging of 55 different loci, which shows that ExTaSy can be used to tag genes across the *Drosophila* genome and demonstrate its versatility for functional studies. We also developed a software that automates guide RNA and homology arm design, aiding efficient synthesis of transgenesis constructs. This novel technology will significantly improve our ability to visualize and manipulate proteins using various applications *in vivo* while maintaining endogenous expression levels and genetic background.

## Main

The ability to visualize and manipulate proteins *in vivo* is central to understanding their roles in development, physiology, and disease. Protein tagging has been instrumental in uncovering fundamental principles of gene regulation, protein dynamics, and cellular organization in *Drosophila*. For example, fluorescently tagged transcription factors have enabled real-time measurements of nuclear dynamics during embryonic patterning, revealing how the concentration-dependent response to Bicoid shapes morphogen gradient interpretation^1^. Quantitative live imaging of endogenously expressed transcription factors has provided insights into chromatin binding dynamics and transcriptional regulation, including the transient association of multiple factors at active regulatory sites^2^. Furthermore, tagging of signaling and adhesion proteins has revealed dynamic cellular mechanisms underlying tissue patterning. For example, fluorescently tagged components of the Notch pathway demonstrated how ligand endocytosis and intracellular trafficking control signal activation during asymmetric cell division and neurogenesis^3^.

Several powerful technologies that enable the analysis of protein function through tagging of genes have been developed in *Drosophila*^4–14^. However, most suffer from drawbacks that limit what conclusions can be drawn from them. Many rely on non-endogenous expression systems, where expression is limited to one protein isoform. Additionally, they increase gene copy number and perturb protein dosage and localization. For example, UAS-driven ectopic expression of transcription factors can lead to cell fate changes^15–18^ while even modest changes in transcription factor dosage can perturb regulatory network function, reflecting the intrinsic dosage sensitivity of gene regulatory systems^19^. BAC- or fosmid-based transgenes, while capturing larger regulatory regions, sometimes fail to reproduce the full spatiotemporal expression patterns of endogenous genes due to incomplete enhancer coverage and sensitivity to chromatin context^20^. Although methods exist for tagging endogenous loci, transposon-mediated strategies insert constructs randomly throughout the genome^4,8,11,12^, thereby lacking the accuracy to precisely modify a target locus in the desired way. The most promising current method using a targeted approach to tag *Drosophila* genes endogenously is CRIMIC (CRISPR-Mediated Integration Cassette)^6,7,10^, where a cassette containing a splice acceptor site and a tag is inserted into an intron between coding exons using CRISPR/Cas9. Depending on the locus structure and integration site, it can be utilized to tag all or select protein isoforms generated by the gene. Flanking *attP* recombination sites in CRIMIC constructs as well as in its randomly inserting predecessor, MiMIC (Minos Mediated Integration Cassette)^14^, allow for an exchange of the cassette and, thus, the primary T2A-GAL4^21^ tag, *via* recombinase-mediated cassette exchange (RMCE) ^22–27^. However, the design features of MiMIC/CRIMIC lead to a truncated tagged protein or tagging within the coding sequence, both of which may affect gene function. Additionally, the insertion of the MiMIC/CRIMIC cassette into a non-coding locus can disrupt regulatory elements and may have undesired effects on the expression of the tagged and/or neighboring genes. The use of a splice acceptor site makes MiMIC/CRIMIC also unsuitable for the tagging of single-exon genes, and while methods exist to circumvent this limitation^7,28^, they rely on non-exchangeable tags or fully replace the coding sequence, limiting downstream applications. Thus, a new tagging strategy which recapitulates endogenous gene expression, is suitable for all genes, leads to minimal modification of the tagged locus, and allows flexibility depending on the desired application would alleviate these drawbacks and therefore be beneficial to the research community.

Here, we present the Exchangeable Tagging System (ExTaSy), a novel CRISPR-based approach for inserting a tag at endogenous coding sequences. ExTaSy utilizes a streamlined single-vector homology-directed repair (HDR) system, allowing for cost-effective and rapid vector construction *via* Gibson assembly with just two synthetic DNA fragments, which can be designed using our automated software. Our tagging vector includes design features from several well-established techniques into a single platform: a visible selection marker, flanked by piggyBac transposon ends for near-scarless excision^29,30^, and asymmetric *attP* and *attB* sites that allow *in vivo* tag swapping through directional ΦC31-mediated recombination^12,22,31^. We showcase the functionality and versatility of ExTaSy in a sample set of genes, most of which encode transcription factors essential for correct development. We demonstrate efficient tagging, marker excision with minimal scarring, and reliable tag exchange. ExTaSy provides a versatile platform for dynamic protein studies in multiple tissues and developmental stages with minimal disruption to the endogenous gene locus.

## Results

### The Exchangeable Tagging System allows for streamlined cloning and efficient genome editing

We designed the ExTaSy plasmids so that the cassettes for HDR and for guide RNA (gRNA) expression are combined in a single molecule (Fig. 1). This enables a streamlined cloning and injection procedure for transgenesis with a single DNA preparation. Separate vectors are used for tagging at the carboxy- and amino-terminus (pExTaSy-EC and pExTaSy-EN, respectively; Fig. 1A, B). Cloning requires the insertion of two synthetic DNA fragments containing left and right homology arm sequences (Fig. S1). The gene-specific gRNA sequence is incorporated at the 5’ end of the left homology arm. To streamline the *in silico* design of these fragments, we developed a bioinformatic pipeline (see Methods and Data Availability) to predict optimal sequences for gRNA, homology arms, and a set of primers for insertion validation for every N- and C-terminus of a user-defined gene list.

**Figure 1:**
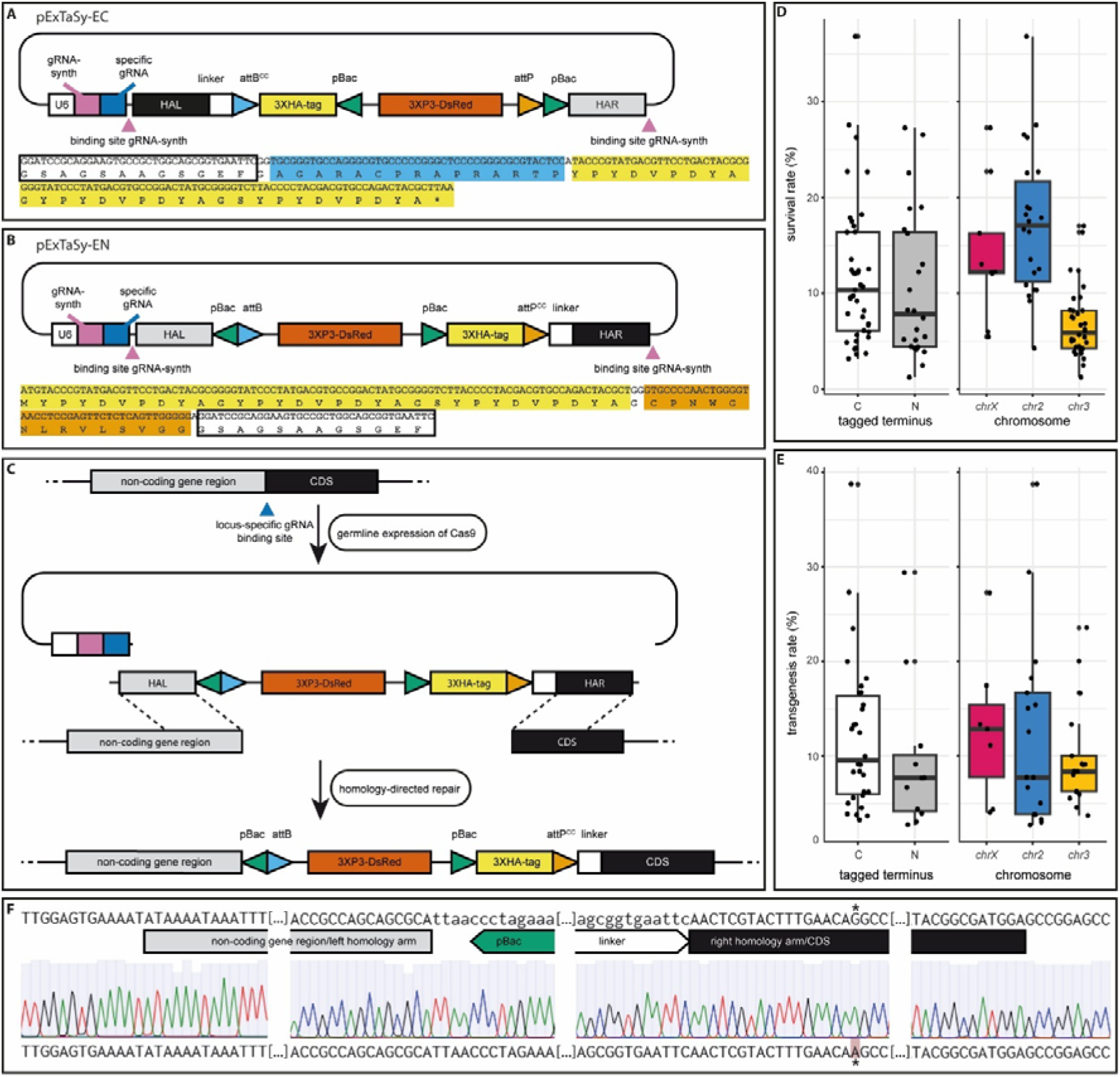
Transgenesis using ExTaSy. (A, B) Schematics of pExTaSy-EC and pExTaSy-EN for C- or N-terminal tagging, respectively. Two gRNAs, gRNA-synth and a gene-specific gRNA, are under control of the *U6* promoter. Homology arms (HAL and HAR) flanking the tagging cassette are used during HDR. The 3XHA tag is connected *via* a linker peptide and an *attB* (pExTaSy-EC) or *attP* (pExTaSyEN) site to the coding sequence (CDS) in the homology arm. The recombination site in the linking *att* has been mutated to CC (*attB*^*CC*^, *attP*^*CC*^). The transgenic marker (3XP3-DsRed) is flanked by piggyBac sites (pBac). An *attP* (pExTaSy-EC) or *attB* (pExTaSy-EN) site allows, together with the *attB*^*CC*^/*attP*^*CC*^ site, for RMCE of the tag. Below each map, the sequences (nucleotide and amino acid) for linker, *attB*^*CC*^/*attP*^*CC*^, and 3XHA are shown with similar shading as in the schematic. (C) After injection of the construct into embryos expressing Cas9 in the germline, the gRNAs induce double strand breaks on either side of the HDR template as well as in the vicinity of the gene terminus (shown here for the N-terminus). During HDR, the linearized plasmid serves as a template, leading to an insertion of the tagging cassette into the genome. (D) Survival rates after injection indicate no bias towards tagging at the C- or N-terminus, but survival is markedly lower for tagging of genes on *chr3*. (E) Transgenesis rates are slightly lower for N-than for C-terminal constructs, but there does not appear to be a strong chromosomal bias. (F) Screenshot of Sanger sequencing analysis after engineering of the *Ultrabithorax* N-terminus. Shown are the 5’ and 3’ ends of the left and right homology arms. A silent mutation (indicated by an asterisk) has been introduced into the CDS to avoid Cas9 cutting the pExTaSy plasmid – see Methods for details.

The homology arms in an ExTaSy plasmid flank the HDR cassette, which contains the primary protein tag [the coding sequence for three haemagglutinin epitopes (3XHA)] and a transgenic marker (3XP3-DsRed; see Fig. 1). The marker is flanked by piggyBac transposon ends that allow for near-scarless removal (see below). The whole construct is flanked by *attP* and *attB* recombination sites, which allow for a subsequent unidirectional tag exchange. One of the sites (*attB* in pExTaSy-EC, *attP* in pExTaSy-EN) connects the tag to a linker peptide consisting of the sequence GSAGSAAGSGEF^32^. Importantly, this linking *attP*/*attB* sequence has been modified so that its recombination site (normally TT) is replaced by CC (*attP*^*CC*^ and *attB*^*CC*^), which prevents recombination between the two endogenous sites^33^.

Translation of the *attP*^*CC*^/*attB*^*CC*^ site extends the linker by 16 amino acids (GAGARACPRAPRARTP) in pExTaSy-EC and 15 amino acids (GCPNWGNLRVLSVGG) in pExTaSy-EN (Fig. 1A, B).

A gRNA expression cassette from pCFD5^34^ lies 5’ of the HDR cassette and contains sequences for two gRNAs, gRNA-synth and a gene-specific gRNA, under control of the *Drosophila U6* promoter. gRNA-synth facilitates Cas9-mediated cutting of the plasmid outside of the homology arms (Fig. 1A), while having no homology to a sequence in the *Drosophila melanogaster* reference genome. Linearization of the HDR template increases HDR efficiency while minimizing homology arm length requirements^35,36^, and a similar strategy has, for example, been used for CRIMIC insertions^6,7^. Simultaneously, the gene-specific gRNA induces a double strand break in the vicinity of the start or stop codon of the targeted gene (Fig. 1C). The linearized plasmid is then used as a template for HDR, which inserts the tagging cassette into the genome. During this process, the native start or stop codon is replaced by the start or stop codon of the 3XHA tag for N- and C-terminal tagging, respectively.

We injected 65 constructs targeting loci on different chromosomes (Fig. 1D, E; Supplementary File S1), and 41 yielded transgenic lines with DsRed marker expression (63.1 % overall success rate). An additional 17 lines were established after injection by BestGene Inc. Correct insertions were confirmed for 44 lines with PCRs that span the 5’ and 3’ ends of the tagging cassette and the corresponding end of the genomic locus, including the full length of the homology arms (Fig. 1F). In-house survival rates after injection were variable and ranged from 1.2 to 36.8 % (average 11.4 %; Fig. 1D). While we could not detect a marked difference for survival rates between N- and C-terminal constructs (average 10.6 and 11.9 %, respectively), constructs inserting into different chromosomes showed varying survival rates: 14.1 % for the X chromosome (*chrX*), 17.3 % for the second (*chr2*), and 6.8 % for the third (*chr3*). The low survival rate for *chr3* injections is likely because the Cas9-expressing construct of the injection line (BDSC_78781)^37^ is homozygous lethal and, hence, kept over a balancer chromosome, which means that only half of the embryos survive whether injected or not.

Transgenesis efficiencies (i.e., number of surviving G_0_ that yielded transgenic offspring) ranged between 2.7 and 38.7 % (average 11.4 %; Fig. 1E). The rate was 12.1 and 9.6 % for C-terminal and N-terminal constructs, respectively. Constructs inserting on *chrX* had a success rate of 12.9 %, those on chr2 one of 12.3 %, and those on chr3 one of 9.9 % indicating that there is no strong chromosomal bias.

While our gRNA selection pipeline is aimed at minimizing the likelihood of off-target effects, we cannot exclude the possibility of mutations induced elsewhere in the genome through non-homologous end joining-based repair of Cas9-induced double strand breaks. However, we have not noticed any instances of the DsRed marker segregating with a different chromosome than the one we targeted, arguing against off-target insertion of our tagging cassette.

### Straightforward marker removal using piggyBac transposase leads to minimal locus modification

While the inserted 3XHA tag is comparatively small (58 amino acids, including linker, *attB*^*CC*^/ *attP*^*CC*^ site, and tag), the whole inserted cassette including the transgenic marker comprises of 1,919 bp for C-terminal and 1,922 bp for N-terminal tagging. Such a large insertion can have negative effects on the expression of the tagged gene. Most genes we tagged at the C-terminus were homozygous viable, indicating limited detrimental effects. However, some lines remained in a heterozygous state, indicating that the endogenous 3’ untranslated region (UTR) and/or the C-terminus of the protein contain essential elements for normal gene function. This is, for example, the case for *Ultrabithorax* (*Ubx*), which is post-transcriptionally regulated through microRNA binding to the 3’UTR^38^. Insertions at the N-terminus always disrupt gene expression because the promoter is displaced from the CDS (Fig. 1C and 2A). Thus, most loci targeted with an N-terminal insertion remained heterozygous. Interestingly however, we noted some exceptions where such lines are homozygous viable, including for *scalloped*. This gene harbors multiple START codons in different exons, and it could be that disruption of the expression of one isoform is not sufficient to abrogate gene function. To re-establish normal gene function, we included flanking piggyBac transposon ends, which enable removal of the 3XP3-DsRed marker through expression of piggyBac Transposase^29,30^. This leaves a minimal scar (TTAA) in front of the tag for N-terminal and after the tag for C-terminal constructs (Fig. 2A, B).

**Figure 2:**
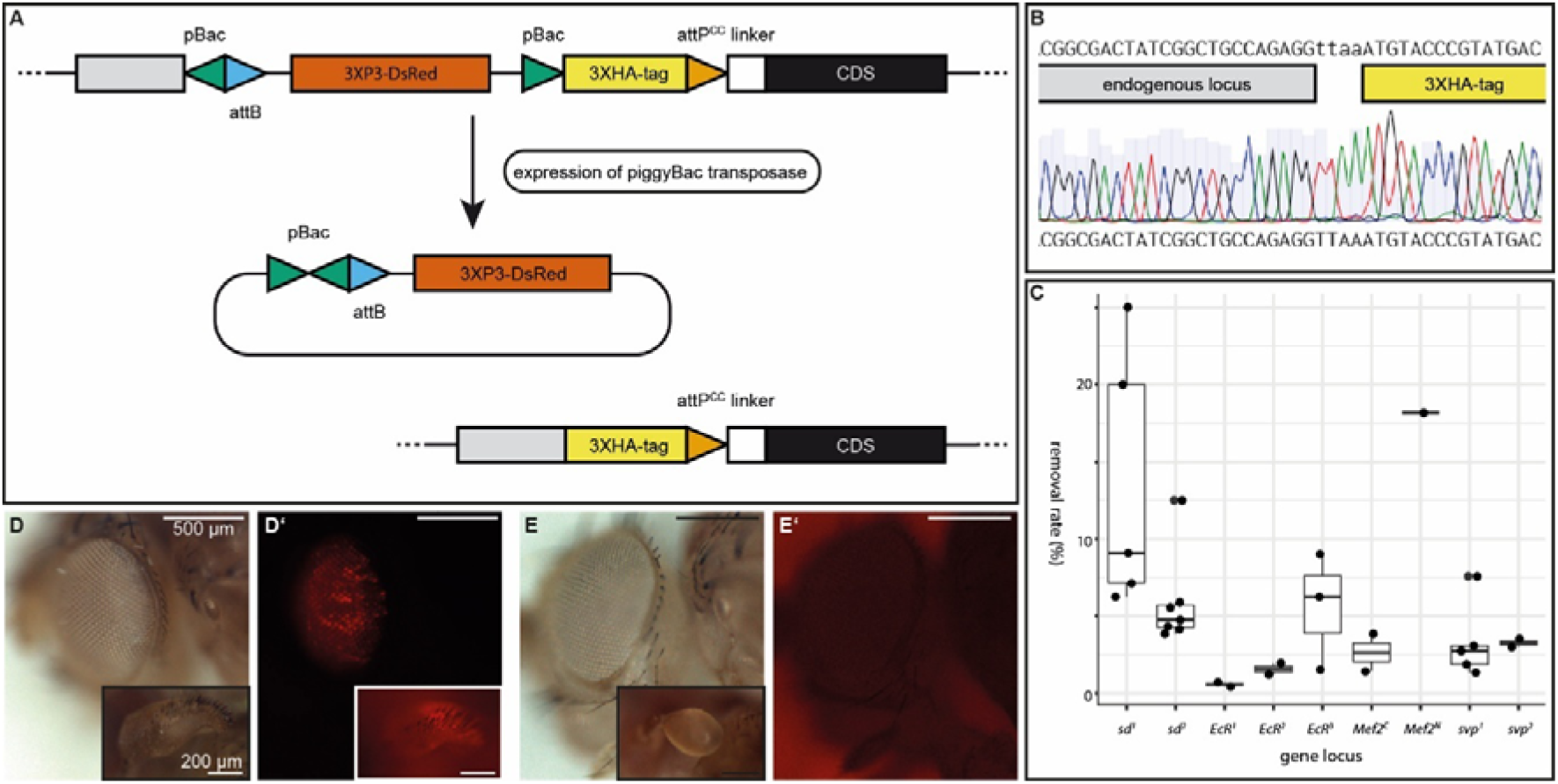
Removal of the transgenic marker using piggyBac Transposase. (A) Schematic of the removal process, shown here for an N-terminal tag. Ubiquitously expressed piggyBac Transposase recognizes the piggyBac transposon ends (pBac) on either side of the transgenic marker cassette and catalyzes their excision. After marker removal, only the tag and linker remain at the locus, as well as a minimal TTAA scar upstream of the tag. (B) Screenshot of Sanger sequencing analysis after marker removal at one of the N-termini of the *Ecdysone receptor* gene (*EcR*) shows the TTAA scar upstream of the 3XHA tag. (C) Success rates of marker removal for different termini of the genes, *scalloped* (*sd*), *EcR, Myocyte enhancer factor 2* (*Mef2*), and *seven up* (*svp*). We define the success rate as percentage of screened flies per cross that yielded offspring with loss of DsRed expression. (D, D’) Flies in which the CDS of *Ubx* has been tagged at the N-terminus with pExTaSy-EN show patchy DsRed expression in the eyes. The inlay shows a haltere that exhibits a partial transformation into a wing. DsRed expression from the transgenic marker is visible in the haltere (inlay in D’). (E, E’) After removal of the transgenic marker cassette, the tagged chromosome no longer requires balancing (i.e., flies become homozygous for the tagged allele), no DsRed expression is detectable, and halteres show a wild-type morphology. Note that the image in E’ has been taken with a much higher gain than D’, leading to a red background.

To test the ability to remove the transgenic marker, we crossed flies harboring an ExTaSy insertion to flies ubiquitously expressing piggyBac transposase (see Fig. 2A and S2). F_1_ offspring were then individually crossed to a balancer line and F_2_ offspring screened for loss of DsRed expression. This was done for 19 independent ExTaSy insertions throughout the genome, 17 of which yielded F_2_ offspring lacking DsRed expression. The two crosses that did not lead to piggyBac removal had a very limited number of F_1_ offspring due to the generally low survival rate of the used piggyBac transposase-expressing fly line. The efficiency of marker removal was quantified for 9 of the 17 insertions (Supplementary File S2). The rate of F_1_ crosses with DsRed-negative offspring varied depending on the locus containing the insertion, ranging between 16.7 and 100 % (average 44.5 %). The success rate within a cross (*i*.*e*., number of negative per total offspring of an individual F_1_) ranged between 0.4 and 25 % (average 6.1 %; Fig. 2C, Supplementary File S2). We performed Sanger sequencing for one established line for each of the 17 loci and confirmed clean excision of the piggyBac sites in all cases (Fig. 2B).

After removal of the marker at the N-terminus, we observed that previously heterozygous lines became homozygous for the tagged allele, indicating that marker removal usually restores normal protein function. This is, for example, the case for *Ubx*, where the loss of function induced by the N-terminal tag is identifiable by a partial haltere-to-wing transformation when kept over a *TM3, Sb*^*1*^ balancer chromosome, which carries a hypomorphic *Ubx* mutation (Fig. 2D). Flies in which the marker was removed developed normal halteres (Fig. 2E) and the allele became homozygous-viable. Additionally, expression of the 3XHA tag became detectable by immunostaining (see Fig. 3). However, in the case of *Mef2*, removal of the transgenic marker did not restore full gene function and flies remained in a balanced state, indicating that the protein terminus is essential for full function, consistent with previous unpublished findings (Hubbert & Taylor, personal communication).

**Figure 3:**
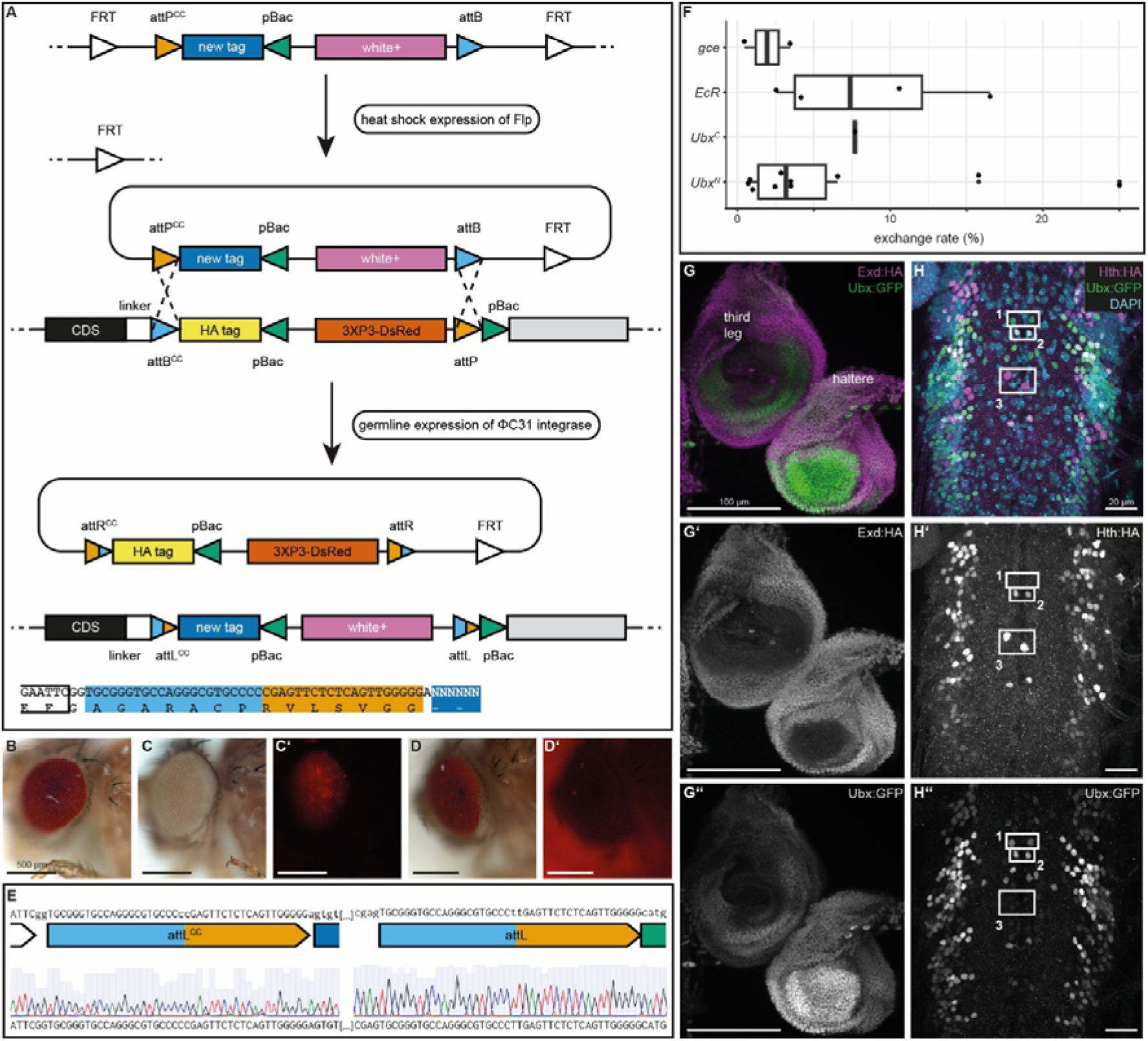
Tag exchange through RMCE. (A) Schematic of tag exchange for the C-terminus. The SwapSy construct is flanked by FRT sites. Flippase induces remobilization, leaving a single FRT site in the genome. ΦC31 integrase catalyzes recombination between the *attP*^*CC*^/*attB*^*CC*^ and *attB*/*attP* sites in the SwapSy and ExTaSy constructs. The exchange of tags and markers is unidirectional and leads to the formation of *attL* sites. The *attL*^*CC*^ site links the new tag to the linker (nucleotide and amino acid sequences indicated below). (B) Flies carrying an sfGFP swap construct have red eyes due to *w*^*+*^ marker. (C, C’) Flies with pExTaSy-EC-tagged *Ubx* express DsRed. (D, D’) Flies in which 3XHA has been exchanged for sfGFP have red eyes and no DsRed expression. The red background in D’ is due to a higher gain than in C’. (E) Snapshot of Sanger sequencing after tag exchange shows the *attL* sites at the *Ubx* C-terminus (*i*.*e*., from flies as in D). (F) Success rates for tag exchange at the C-terminus of *germ cell-expressed bHLH-PAS* (*gce*), *EcR*, and *Ubx*, as well as the N-terminus of *Ubx*. (G-G’’’) Immunostaining of HA-tagged Exd (magenta) and sfGFP-tagged Ubx (green) in leg and haltere imaginal discs. Exd exhibits uniform expression, but no expression in central regions (G, G’). Ubx shows strong expression in central and ventral regions and weaker expression in the remaining haltere disc cells (G, G’’). In third leg discs, Ubx is expressed in ventral regions of the developing coxa and femur (G, G’’). (H,H’’’) Immunostaining of HA-tagged Hth (magenta) and sfGFP-tagged Ubx (green) in the ventral nerve chord. Anterior is to the top. DAPI staining in cyan shows nuclei. Neurons in box 1 only express Ubx, neurons in box 2 express both Ubx and Hth, and neurons in box 3 only express Hth.

### Tag swapping through recombination-mediated cassette exchange (RMCE) enables various downstream analyses

While the 3XHA tag can be used for applications like immunostaining and immunoprecipitation, one might also wish to examine protein function in other ways, such as targeted degradation or the discovery of interaction partners. The true power of ExTaSy lies in the fact that the primary tag can be exchanged for any other using RMCE^22,39^. This is possible through three rounds of fly crosses (Fig. S3), making the system more accessible for most researchers than having to create transgenic flies with different tags for each application through injections, which is laborious and requires a specialist set-up. Additionally, the genetic background of flies with swapped tags can be kept the same through balancer crosses, making experiments easily comparable.

The process of tag exchange requires three components on different chromosomes to be combined in one fly (Fig. 3A, Fig. S3): (1) an ExTaSy construct with primary 3XHA tag, (2) a donor construct with the new tag, and (3) the enzymes needed for remobilization of the donor construct (Flippase recombinase) and RMCE (ΦC31 integrase)^40,41^. We have engineered swap constructs for C- and N-terminal tag exchange, pSwapSy-C and pSwapSy-N, respectively (Fig. 3A, Fig. S4). The swap cassette consists of a new tag and a new transgenic marker, a *white* rescue construct (*w*^*+*^), which is, like 3XP3-DsRed the ExTaSy construct, flanked by piggyBac sites for near-scarless removal. The swap cassette is flanked by *attB*/*attP* sites to allow unidirectional recombination between them and the corresponding sites in the original ExTaSy construct. The whole construct is flanked by Flippase recombinase target (FRT) sites. Using CRISPR/Cas9, we have created donor lines carrying SwapSy constructs with sfGFP as the new tag at defined landing sites on *chr2* or *chr3*^42^ (Fig. 3B).

In flies carrying all three tag exchange components, heat shock is used to ubiquitously express Flippase, which mediates remobilization of the swap construct *via* recombination at the flanking FRT sites (Fig. 3A). ΦC31 integrase expression is limited to germline cells, where it can catalyze the recombination between *att* sites in the SwapSy and ExTaSy constructs. This results in the creation of *attL* sites at either end of tagged gene locus, one of which now connects the tag to the linker peptide and translates into GAGARACPRVLSVGG (Fig. 3A, Fig. S3). Instances of tag exchange are identified in the next generation by screening for red eye color and selecting against DsRed expression (Fig. 3C-E). We have attempted to exchange the 3XHA tag for sfGFP at five loci, and the only case where this was unsuccessful was the C-terminus of *Methoprene tolerant* (*Met*). The rate of single crosses per locus that yielded positive swaps ranged from 10.5 to 25 % (average 15.6 %), and the ratio of positive flies within a single cross from 0.5 to 25 % (average 6.7 %; Fig. 3F, Supplementary File S3). We used Sanger sequencing to confirm the tag exchanges and found one line out of four for the *Ubx* N-terminus where the formation of the *attL*^*CC*^ site resulted in the insertion of an additional 9 bp (CGGCGTGCT) at the recombination site. All other analyzed lines showed the expected formation of *attL* sites and integration in the anticipated orientation, indicating that the configuration of the *att* sites surrounding the tag indeed leads to unidirectional integration.

We compared the expression of both tags at *Ubx* by crossing a C-terminally tagged *Ubx*^*swap-sfGFP*^ line to a 3XHA-tagged line in which we removed the N-terminal transgenic marker and performing immunostaining on haltere and leg imaginal discs (Fig. S5A). Expression of the two tagged alleles overlaps spatially in apparently all cells. It also largely overlaps in the larval ventral nerve chord; however, few cells exhibited stronger staining for GFP-tagged Ubx (Fig. S5B). This may be because this allele does not express the native *Ubx* 3’UTR due to the presence of the transgenic marker 3’ of the tag CDS and may, thus, lack sequences for correct translational control.

We also crossed the *Ubx*^*swap-sfGFP*^ line to flies in which Ubx co-factors, Extradenticle (Exd) and Homothorax (Hth), were tagged at the C-terminus. This allows for co-staining of both proteins and an analysis of where the two transcription factors may interact to regulate target gene expression. Examining haltere and third leg imaginal discs (Fig. 3G), we found that Exd and Ubx are expressed similar to published patterns^43–45^. Exd is found uniformly throughout both discs, but not in the center, which will develop into the distal parts of each appendage. Ubx shows strong expression in the center and most ventral region of the haltere disc and weaker expression throughout the disc. Ubx expression in the third leg disc appears to be limited to the ventral side of the area developing into coxa and femur of the adult leg. Thus, Ubx and Exd overlap in parts of both discs but also show unique expression patterns. Ubx and Hth were compared in the ventral nerve chord (Fig. 3H), where expression of both proteins is complex and varies in strength between different neurons^46^. We found cells that co-express both proteins, but also neurons that only express either Ubx or Hth. We also found cells in the periphery of the nerve chord which express both transcription factors, but only Ubx is localized in the nucleus while Hth remains cytoplasmic (Fig. S5C). Note that the tag we introduced into *hth* only tags isoform E (FBpp0099878), and other protein variants of Hth may show different expression patterns.

## Discussion

We have shown that ExTaSy can faithfully tag proteins in *Drosophila* with 3XHA with the option of tag exchange and transgenesis marker removal. Our survival rates after injection were relatively low for some constructs. However, average rates are comparable to published values for HDR-based genome editing strategies^47,48^. We outsourced injections of fifteen additional constructs and also some of these induced high mortality. We speculate that, even though Cas9 expression should be germline-restricted^48,49^, somatic CRISPR-induced mutations could lead to increased mortality when targeting certain gene loci.

We confirmed tagging cassette insertion at the correct genomic locus using breakpoint-spanning PCRs, a standard approach in the field (*e*.*g*., ^6,7,10^). While highly specific, these PCRs may miss head-to-tail insertions or other structural variations reported in mammalian genome engineering^50,51^. Whole-insert-spanning PCRs could address this, but are more effective in homozygous flies, as the untagged allele is otherwise preferentially amplified.

There are several advantages of our system over other transgenesis procedures currently used in *Drosophila* (see Table 1). The one-vector approach allows for streamlined and cost-effective cloning of the tagging plasmid, whereas most CRISPR-based systems require co-injection of separate plasmids for gRNA expression and the HDR cassette. ExTaSy thus halves the time and cost of cloning.

**Table 1:**
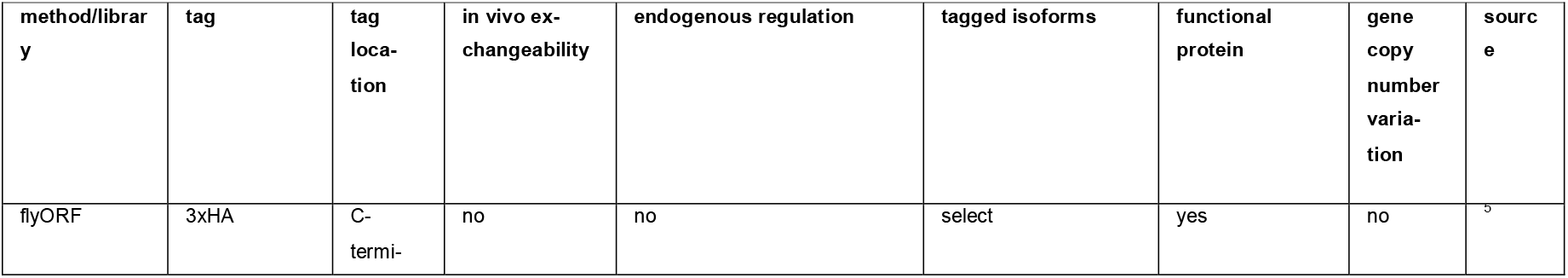
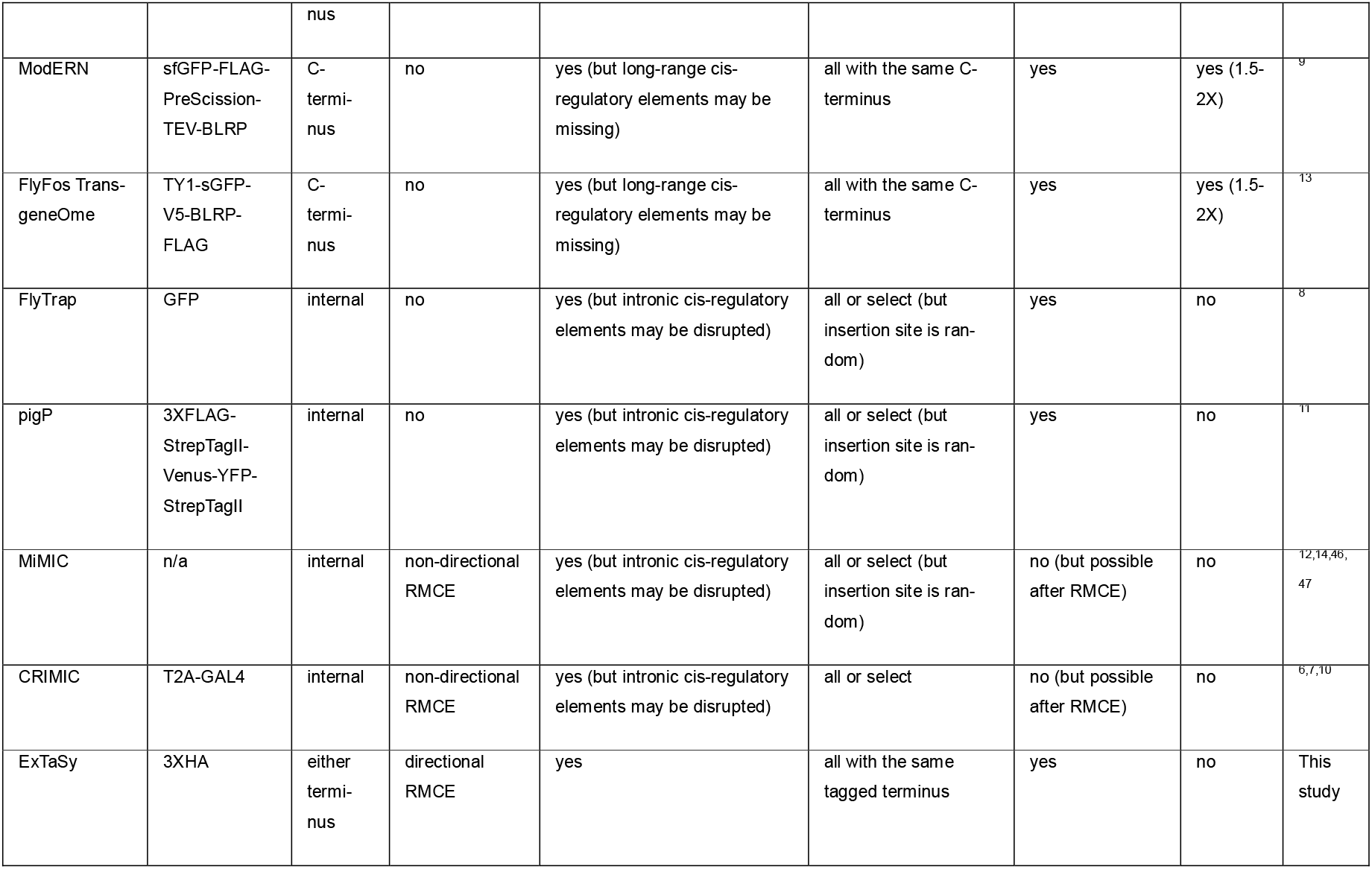
Comparison of different strategies for protein tagging in *Drosophila*.

Additionally, since the tag can be swapped using fly crosses, transgenic lines need only be created once even if a different tag is desired at a later point. Flies with a new tag can then be generated even by research groups lacking the expertise or means of creating transgenic lines through injection. While MiMIC^12,14,52,53^ and CRIMIC^6,7,10^ also allow tag exchange via RMCE, their use of inverted *attP* sites makes the process non-directional^22,33^, increasing the number of lines that must be screened. In contrast to other RMCE systems, the *att* site configuration in ExTaSy results in unidirectional swapping of tags^33^. This removes the need to screen for swaps with the correct orientation, increasing the efficiency of the system. However, since the tag swap takes place 3’ of the start or 5’ of the stop of the coding sequence (for N-terminal and C-terminal tags, respectively), the *att* site is translated and incorporated into the protein between the tag and a linker peptide^32^. While we have selected an open reading frame translating the connecting *att* site into an amino acid sequence that as closely as possible reflects that of the linker, we cannot categorically exclude a negative influence of the *att* site translation on the tagged protein or the tag itself. However, we were able to detect the HA tag using immunostaining of C-as well as N-terminal tags, indicating that the tag works as expected, and since transgenic lines often become homozygous, generally, no major implications for the tagged protein itself are expected.

Using ExTaSy, tags are introduced at the endogenous genomic locus, which keeps gene regulation intact and allows for the individual tagging of alternative protein isoforms if they differ in their terminus. Existing resources to analyze tagged proteins in *Drosophila* often rely on induced expression (*e*.*g*., FlyORF^5^), altering the protein level and/or potentially inducing expression in additional cell types, tissues, and organs. Moreover, individual transgenic lines are required for each protein isoform. Methods using BACs or fosmids to insert an additional tagged gene copy in a defined landing site (e.g., ModERN^9^ and FlyFos TransgeOme^13^) may recapitulate endogenous expression but can lack long-range *cis*-regulatory elements. Because the endogenous allele is unmodified, they also lead to copy number variation that may cause higher expression levels of the analyzed protein, and the untagged protein cannot be manipulated. Other resources for tagging genes endogenously (FlyTrap^8^, pigP^11^, MiMIC^12,14,52,53^, and CRIMIC^6,7,10^) insert into introns that separate coding exons and utilize splice acceptor sites to tag all or select isoforms, depending on the insertion site. However, only CRIMIC is inserted site-specifically *via* CRISPR, while MiMIC, pigP, and FlyTrap use random transposon-based insertion. The insertion into non-coding gene sequences could disrupt *cis*-regulatory elements and have a negative effect on the expression of the tagged and/or neighboring genes. ExTaSy avoids these issues by targeting the endogenous locus between the coding sequence and UTR of a gene, minimizing disruption of *cis*-regulatory elements and maintaining endogenous regulation.

The availability of separate ExTaSy vectors for the C- and N-terminus allows protein analysis even if the tag at one of them leads to non-functionality. N-terminal tagging is possible because gene function can be restored after near-scarless marker removal with piggyBac sites, a method pioneered in this way by the O’Connor-Giles lab (unpublished data and ^29^). Marker removal is also advisable for C-terminal tags since the 3xP3 sequence commonly used to drive transgenic marker expression in *Drosophila* can lead to ectopic expression of the tagged or other nearby genes^54^. While we found that, generally, tagged alleles become homozygous after marker removal, indicating that normal gene function is restored, it must be considered that the remaining TTAA scar in the Kozak sequence could reduce protein levels. Indeed, a study testing functional effects of changes to the Kozak sequence found that a mutation of the canonical CACCatgA to TTAAatgA reduces activity to about 79 %^55^. This sequence closely resembles the one in our system (TTAAatgN), and a similar drop in activity may occur, although the canonical sequence is only displaced in our case. Additionally, the TTAA scar may also have a negative effect on the 3’UTR. This must be evaluated on a gene-by-gene basis. The genes that we tagged for this study largely encode transcription factors whose function is indispensable for normal development, emphasizing that the TTAA scar has a minor effect on gene expression.

Recently, methods have been developed in flies that allow for a truly scarless removal of transgenesis markers using CRISPR/Cas9. The SEED/Harvest system^56^ utilizes a SEED (scarless editing by element deletion) donor that contains a transgenic marker flanked by “Harvest” gRNA sites. Cas9-induced double strand breaks on either side of the SEED insert lead to its removal and are repaired through micro-homology end joining between very short homology arms in the remaining transgenic insert, making the tag only functional after marker removal. Modifying our ExTaSy constructs to include a SEED cassette would leave *attP* and *attB* (or, after tag exchange, *attL*) sites at either end of the inserted construct that would have to be incorporated into the coding sequence to avoid them leaving a scar. The cotranslation of the linking *att* site in our ExTaSy constructs does not appear to affect the functionality of the tag or the tagged protein, but the presence of an additional site at the other end of the tag would require further testing. A construct design with linking *att* sites at either end would also allow internal rather than terminal tagging, which may be advantageous where both C- and N-terminal tags are non-functional, fail to capture all isoforms, or to allow the manipulation of specific isoforms generated through alternative splicing. Tools and resources for the prediction of protein structure^57,58^ and interaction domains^59^ could be integrated into our bioinformatic pipeline to aid optimal insertion site selection. Conceivable ExTaSy constructs for internal protein tagging would then include linkers at either end of the tagging cassette that flank the *attP* and *attB* sites as well as a SEED/Harvest-like removable transgenic marker, since, in this case, a remaining TTAA scar would induce a frameshift mutation.

The possibility of tag exchange makes ExTaSy highly versatile. This is similar to MiMIC^12,14,52,53^, and CRIMIC^6,7,10^, where RMCE can be used to exchange the initially inserted construct for a different tag. While the initial tag in MiMIC and CRIMIC lines leads to the expression of a truncated protein that may lack important endogenous functions and requires a tag exchange before the protein itself can be analyzed, the co-expression of GAL4 via the T2A system^21^ allows for other types of experiments. Conversely, 3XHA-tagged ExTaSy lines can in principle be used immediately for experiments in which a fully functional protein is detected with antibodies. In rare cases, this may be disadvantageous. It is conceivable that the inclusion of a tag at the C-terminus of an otherwise functional protein may lead to faulty post-translational regulation and the creation of a dominant mutation. It would not be possible to create stable transgenic lines for thus affected genes. This could be alleviated by creating ExTaSy constructs that use T2A-GAL4 as initial tag, with the above-mentioned limitations. We have shown that different tags can be used for co-staining of different proteins in the same tissues; however, the sfGFP tag in our SwapSy constructs is also useful for targeted protein depletion using the deGradFP system^60^ or live imaging if the expression level is high enough. Additional swap donor constructs can easily be generated using our open platform, where cloning of novel tags into our SwapSy constructs is straightforward. This will enable researchers to analyze protein function in various ways while maintaining the same genetic background through controlled crosses, making results highly replicable. Potential tags include inducible degrons^61,62^ for loss-of-function studies, tags that are recognized by nanobodies (*e*.*g*., ALFA^63^), biotin ligases (*e*.*g*., TurboID^64^) to study the interactome *via* proximity labeling, and Llama tags^65^ for *in vivo* visualization of protein dynamics. Moreover, co-translation of GAL4 or fluorescent proteins using the T2A system^21^ will make it possible to label cells by the expression of a protein or individual isoform that will be expressed without a tag and, hence, be completely functional.

While we have shown here the tagging of different transcription factors, ExTaSy serves as a broadly applicable platform allowing analysis of virtually any *Drosophila* protein without the need to generate specific antibodies. This makes the system also interesting for use in other animals. Our bioinformatic pipeline can, in principle, be adapted to design these sequences for any species if the genome sequence and gene annotations are provided. However, some specialized input files (*e*.*g*., a pre-existing database of gRNAs with off-target predictions) are only available for *Drosophila*, which requires further modification of the pipeline. Additionally, some of the functional features of ExTaSy like marker removal and tag exchange will be more difficult to implement in other species. Yet, the use of a single plasmid system for transgenesis is appealing and will allow at least C-terminal tagging of genes. This may require the exchange of the *U6* promoter and the promoter in the transgenesis marker for species-specific sequences^66^. We hope that widespread adoption of ExTaSy by the research community will enable novel insights into protein function in *Drosophila* as well as comparative studies across species.

## Methods

### *Drosophila* lines and husbandry

Flies were kept on standard cornmeal/agar food at 18 °C with a light/dark cycle of 12:12 hours. During experiments, expansion of stocks, crosses, *etc*., flies were kept at 25 °C with a light/dark cycle of 12:12 hours. Table S1 shows the *Drosophila* lines used in this study.

### Bioinformatic pipeline for sequence design

A detailed description is available in our GitHub repository (https://github.com/emmacwatts/AutoTagsCRISPR/tree/main). In short, gRNAs that sit directly at the start or stop codon in a way that separates the protospacer-adjacent motif (PAM) and the gRNA binding site are preferred. If unavailable, gRNAs are filtered for having a 3’ overhang of less than 12 bp in a homology arm, after which the one that cuts closest to the start/ stop is selected. This approach avoids recognition of the HDR plasmid by the gRNA^47^. If only gRNA binding sites with 12 or more bp overhang are present, the one located closest to the start/stop codon within the coding sequence (CDS) is selected. Since the presence of the gRNA and PAM sequence in the homology arm would induce cutting of the HDR template, synonymous mutations are introduced in the HDR plasmid. Where possible, one of the Gs in the PAM (see Fig. 1E) is mutated. Alternatively, two nucleotides in the 3’ region of the gRNA binding site are mutated, which leads to a lower likelihood of Cas9 binding^47,48^. In addition to gRNA and homology arm selection, a set of primers for validation of successful tagging is designed using Primer3^67,68^ based on a set of stringency parameters of GC content, melting temperature, and position to optimize for ideal conditions.

### Molecular cloning

Creation of vector maps and alignment after Sanger and whole-plasmid sequencing was done with Benchling (https://benchling.com). All Restriction enzymes were from New England Biolabs (NEB) and used according to the manufacturer’s guidelines. PCRs to generate fragments for cloning were carried out using Q5® High-Fidelity 2X Master Mix (NEB #M0492). NEBuilder® HiFi DNA Assembly Master Mix (NEB #E2621) was used for Gibson assemblies according to the manufacturer’s instructions. E. coli DH5α cells (grown in house) were used in a standard heat-shock transformation protocol, using ampicillin selection for all constructs. Correct clones were confirmed with colony PCR using OneTaq® Quick-Load® 2X Master Mix with Standard Buffer (NEB #M0486). Synthetic DNA fragments were ordered from Integrated DNA Technologies (IDT) or Twist Bioscience, for sequences see Table S2. Primers were ordered from IDT, for sequences see Table S3. Plasmid DNA was harvested using either the Monarch Plasmid Miniprep kit (NEB #T1110) or the Purelink HiPure Midiprep kit (ThermoFisher #K210005). Constructs were verified by Sanger sequencing, carried out by Source BioScience or Eurofins Genomics.

#### pExTaSy constructs

To create pExTaSy-EC and pExTaSy-EN (for tagging at the C- and N-terminus, respectively), a five-fragment Gibson assembly was used. The vector backbone and transgenic marker insert were amplified from pHD-ScarlessDsRed (Kate O’Connor-Giles; Addgene #64703; DGRC #1364) with primers pExTaSy-E_bb_F1 and pExTaSy-E_bb_R1 (backbone), pExTaSy-E_ins_F1 and pExTaSy-E_ins_R1 (insert of pExTaSy-EC), and pExTaSyE_ins_F1 and N-term_DsRed-rev (insert of pExTaSy-EN). The other fragments were synthesized.

#### pSwapSy constructs

pExTaSy-EC was digested with SfoI, SrfI and PvuII, to isolate a 2,336 bp fragment containing an origin of replication and ampicillin resistance gene. This was then used in a four-fragment Gibson assembly reaction, along with two synthesised DNA fragments and a PCR amplicon from pRMCE{G418R-5FCS-GMRWhite}-Vasa-FC31^69^ (Addgene #165887) with primers GMR-w-hsp70-F1 and GMR-w-hsp70-R1 to generate vectors pSwapSyN and pSwapSyC. Antibiotic resistance genes were PCR-amplified from existing plasmids^69^ (BlastR: pBlastR-MCS-QF-MiniWhite, Addgene #165906; neoR: pRMCE{G418R-5FCS-GMRWhite}-Vasa-FC31, Addgene #165887; PuroR: pPuroR-MCS-LexA-VP16-MiniWhite, Addgene #165898) using primers GMR-w-hsp70-F1 and GMR-w-hsp70-R1 and separately ligated into NheI and Acc65I-digested pSwapSyN and pSwapSyC to generate pSwapSyN-neo and pSwapSyC-neo, pSwapsyN-Puro and pSwapsyC-Puro, and pSwapsyN-Blast and pSwapsyC-Blast. To allow insertion of the SwapSy constructs into six defined genomic positions on *chr2* and *chr3*^42^, Gibson assembly was used following vector digestion with SapI and PCR-amplification of homology arms using primers with the prefix Zh30, Zh51, Zh58, Zh64, Zh86, and Zh96 (Table S3). Genomic DNA as template for homology arm amplification was derived from injection strains BDSC_78782^37^ for Zh64, Zh86, and Zh96, and BDSC_78781^37^ for Zh30, Zh51, and Zh58 arms. Finally, the CDS for sfGFP was PCR-amplified from pScarlessHD-sfGFP-DsRed (Kate O’Connor-Giles; Addgene #80811) and cloned into the BsaI-digested pSwapSy vectors *via* Gibson assembly.

#### hs-Flp, vas-int constructs

The pattB-yellow body vector (DGRC 1450^28^) was digested with ApaI and SpeI. A synthetic DNA fragment containing a *Drosophila* codon optimized flippase CDS flanked by an hsp70Bb promoter element and an AcNPV p10 3’UTR and an sgRNA-synth1 recognition sequence was digested with ApaI and AvrII and cloned into the digested pattB-yellow body vector to generate pFlpPhi_int1. pFlpPhi_int1 was digested with SnaBI and used in a Gibson assembly reaction with three additional synthetic DNA fragments (vas5’, DmPhiC31 and vas3’) which introduces a *Drosophila* codon-optimized CDS for PhiC31 flanked by a vasa promoter and 3’ region and a second sgRNA-synth1 recognition sequence to generate pFlpPhi_int2. A gypsy insulator element flanked by SpeI and NotI restriction sites was amplified by PCR from the pValium20 vector (DGRC 1467^70^) using primers SpeI-gypsy_F1 and NotI-gypsy_R1, digested and ligated into SpeI-Not1 digested pFlpPhi_int2 to generate pFlpPhi_int3. A second gypsy insulator element flanked by AatII and SacII was amplified from pValium20 with primers AatII-gypsy_F1 and SacII-gypsy_R1, digested and ligated into AatII-SacII digested pFlpPhi_int3 to generate pFlpPhi. Left and Right Homology arms (HAL and HAR) of ∼630 bp for two different genomic locations (zh2A and zh51D) were PCR amplified from genomic DNA from nosCas9-attP2 flies (BDSC_78782^37^) and inserted into SnaBI-PmeI linearized pFlpPhi using Gibson assembly, yielding pFlpPhi_zh2A and pFlpPhi_zh51D. The yellow cassette was removed by digesting pFlpPhi_zh2A and pFlpPhi_zh51D with AgeI and AvrII and a 3xP3_GFP synthetic construct was introduced using Gibson assembly to generate pFlpPhi_GFP_zh2A and pFlpPhi_GFP_zh51D.

#### Gene-specific pExTaSy vectors – see Fig. S1 and Supplementary File S1

Two spacer sequences, one separating the gRNA expression cassette from the HDR cassette, the other at the 3’ end of the HDR cassette, were removed from pExTaSy-EC or pExTaSy-EN by restriction digest with SapI. This generated two DNA fragments of 3,500 and 1,919 bp for pExTaSy-EC and 3,500 and 1,922 bp for pExTaSy-EN, containing the backbone including the gRNA expression cassette, and the HDR cassette, respectively. To clone homology arms and a gene-specific gRNA into the ExTaSy plasmid, a four-fragment Gibson assembly was used. The additional fragments were generated through DNA synthesis by Twist Biosciences or GenScript (Supplementary File S1). The 5’ fragment contained the gene-specific gRNA and gRNA scaffold, 51 bp of the regulatory region downstream of the *Drosophila U6* gene, the binding site for gRNA-synth including the PAM, a 30 bp spacer, and the gene-specific left homology arm for HDR (240 bp upstream of the start or stop codon). Including flanking 30 bp homology sequences for Gibson assembly, this fragment had a size of 500 bp. The 3’ fragment contained the right homology arm (240 bp downstream of the start or stop codon) flanked by 30 bp regions for Gibson Assembly.

### Transgenesis

We largely followed the injection protocol by the Gompel lab (http://gompel.org). Briefly, eggs of Cas9-expressing fly lines (see Table S1) were collected for 30 minutes at 25 °C and lined up on the edge of a cover slip on microscope slide without dechorionation, briefly dried, and overlaid with olive oil. 1.0 mm OD borosilicate capillaries with omega dot fiber (Harvard apparatus #30-0019) were pulled into injection needles using P-1000 Flaming/Brown Micropipette Puller (Sutter Instruments) with a box filament (Sutter Instrument #FB230B) and the following parameters: Pressure: 300, Heat: 505, Pull: 20, Velocity: 60, Delay: n/a, Time: 250, Looping: none. The ramp value was adjusted each time. Needles were filled with transgenesis plasmid diluted with water to 220 ng/µl using microloader tips (Calibre Scientific #930001007) and subsequently opened by breaking them at the edge of the cover slip under the microscope. Injections into the posterior end of the embryos were done with a Femtojet or Femtojet 4i (eppendorf). Approximately 250 embryos were injected per construct. The olive oil was washed off the embryos by briefly rinsing with 100 % ethanol, followed by rinsing with deionised H2O. Cover slips with injected embryos were inserted into fly food in standard vials and kept at 25 °C.

Surviving G0 adult flies were crossed individually to y1, w1118 flies (BDSC_6598). Offspring were selected for expression of DsRed in the case of pExTaSy tagging and for red eyes in the case of swap line generation. Screening for DsRed was largely done at larval and pupal stages. In larvae, expression is detected in eye imaginal discs, central nervous system, and, prominently, anal pads. After pupation, the cells of the anal pads individualize and DsRed expression in the abdomen is still visible for some time as punctae. During later pupal stages, DsRed becomes expressed in the developing eyes. In adults, expression in compound eyes and ocelli is most prominent (see Fig. 2, 3).

Stable transgenic stocks were established through single crosses of transgenic individuals to an appropriate balancer line (see Table S1). Where possible, we used males carrying the insertion in these crosses to avoid meiotic recombination and keep a homogeneous genetic background. In a second round of selection, balanced individuals with loss of the chromosome with the Cas9 transgene were back-crossed to the balancer to establish a stable stock. Injections and selection of DsRed-positive flies for tagging of *Ubx, hth*, and *exd* was done by BestGene Inc (https://www.thebestgene.com).

Correct insertion was confirmed with PCR and Sanger sequencing. Briefly, total DNA was extracted by grinding 1-3 flies using a micropestle in 50 µl squishing buffer (10 mM Tris-HCl pH 8, 1 mM EDTA, 25 mM NaCl, 100 µM Proteinase K). Samples were incubated at 37 °C for 30 minutes, followed by 95 °C for 3 minutes. Samples were briefly centrifuged at 7,000 rpm for 1 minute, and supernatant was removed for use in subsequent PCR reactions. Genotyping PCRs were carried out using OneTaq® Quick-Load® 2X Master Mix with Standard Buffer (NEB #M0486). Two PCRs per inserted fragment were done with primers binding to the 5’ and 3’ ends of the inserted DNA and corresponding gene-specific validation primers – see Table S3 and Supplementary File S1 for details.

### Marker excision and tag exchange

For excision of the 3XP3-DsRed marker, please refer to Fig. S2. Flies with an ExTaSy insertion were crossed to a line carrying on its *CyO* balancer chromosome a transgene that induces ubiquitous expression of piggyBac Transposase under control of an α-Tubulin regulatory sequence (BDSC_8285). Offspring with the ExTaSy insertion (identified by DsRed expression) and piggyBac Transposase were crossed to an appropriate balancer. In the F2 generation, balanced flies with a loss of DsRed expression were selected and crossed again to a balancer to establish a stock. In cases where no flies with a loss of the marker were found, we crossed F2 flies with the ExTaSy insertion and piggyBac Transposase to a balancer and screened the F3 generation for loss of DsRed expression. Loss of 3XP3-DsRed was confirmed with PCR and Sanger sequencing. Genomic DNA was extracted and a genotyping PCR was done as described above (see Table S3 and Supplementary File S1 for primer sequences).

For tag exchange, please refer to Fig. S3. We crossed flies with an ExTaSy insertion to flies with a SwapSy construct on *chr2* or *chr3*. Male offspring were then crossed to a fly line with a transgenic construct on *chrX* (BDSC_33216)^40,41^ or *chr2* (see above) that leads to expression of Flippase Recombinase under control of a heat shock promoter (hs-Flp) and to expression of ΦC31 Integrase in the germ line under control of vasa regulatory sequences (vas-int). Crosses were allowed to lay eggs for two days, then transferred to a new vial. The old vial with developing offspring was placed for 1 h into a water bath at 37 °C. The heat shock was repeated on the next two days. Developing males in which Flp had remobilized the SwapSy construct with its *w*^*+*^ genetic marker from its genomic location were identified by eyes that expressed *w* in a mosaic fashion and showed, thus, red coloration of only some ommatidia of the compound eyes. In germline cells of these flies, ΦC31 Integrase can catalyze the recombination between *attP* and *attB* sites on the original tag construct and the remobilized SwapSy construct (see Fig. 3, Fig. S4). They were crossed to appropriate balancer flies to balance the gene locus at which the exchange of tags may have taken place. Offspring of this cross were selected based on (1) gain of the *w*^*+*^ marker (*i*.*e*., red eyes) and (2) loss of the DsRed marker and crossed individually to a balancer to create a breeding stock. Correct tag exchange events were confirmed with PCR and Sanger sequencing as described above. For primers, see Table S3 and Supplementary File S1.

### Immunostaining and imaging

Larvae were dissected and tissues fixed and stained according to established protocols^71,72^. Briefly, we dissected larvae in ice-cold phosphate-buffered saline (PBS) and fixed tissues for 20 minutes in PBS with ∼4 % formaldehyde. Samples were then washed several times with PBS with 0.1 % Triton-X 100 (PBST) and blocked for 30 minutes in blocking solution (PBST with 5 % normal goat serum and 0.1 % bovine serum albumin). Samples were incubated for ∼16 h at 4 °C in blocking solution with primary antibodies. The following antibodies were used: rabbit-anti-HA (DSHB anti-HA rRb-IgG; dilution 1:300), mouse-anti-HA (DSHB anti-HA rMs-IgG1; dilution 1:300), chicken-anti-GFP (Invitrogen #A10262; dilution: 1:600). Samples were washed at least three times for 10 minutes each with PBST followed by another blocking step for 30 minutes. Secondary antibodies were added in blocking solution and incubated for 2 h at room temperature. The following antibodies were used at a dilution of 1:600: goat-anti-chicken-Alexa488 (Invitrogen #A32931), goat-anti-rabbit-Alexa488 (Invitrogen #A11008), goat-anti-rabbit-Alexa546 (Invitrogen #A11035), goat-anti-mouse-Alexa488 (Abcam #ab150113), goat-anti-mouse-Alexa647 (Invitrogen #A28181). Samples were washed at least three times for 10 minutes each with PBST. DAPI was added in one of the washing steps. Samples were mounted on microscope slides in VECTASHIELD® PLUS Antifade Mounting Medium (Vector Laboratories #H-1900) and imaged on a Zeiss 800 LSM confocal microscope with a 20X lens. Image analysis and manipulation was performed with Fiji^73^.

## Supplementary Files

**Supplementary File S1: Table in TXT format (tab-delimited) containing information on gene-specific constructs and quantifications of transgenesis**. Columns indicate an internal identifier (“id”), symbol (“locus”), CG number (“CG_number”), and FBgn identifier (“FBgn_id”) of the targeted gene, whether C or N-terminus was tagged (“tag_location”), which protein isoforms are covered by the tag (“isoforms_tagged”), targeted chromosome (“chr”), used injection line (“injection_line”), cloned gRNA sequence (“gRNA_sequence”), the offset of the CRISPR cut site relative to the targeted start or stop codon (“gRNA_offset”), the reference genome sequence of the left homology arm (“HAL_refseq”), whether a silent mutation was introduced into the left homology arm (“HAL_mutation”), the ordered synthetic left DNA fragment (“left_fragment”), the reference genome sequence of the right homology arm (“HAR_refseq”), whether a silent mutation was introduced into the right homology arm (“HAR_mutation”), the ordered synthetic right DNA fragment (“right_fragment”), left (“val_primer_F”) and right (“val_primer_R”) validation primer for PCR to confirm transgenesis, number of injected embryos (“n_injected”), number of survivors (i.e., adult flies) after injection (“n_survivors”), rate of survivors per injected embryos (“survival_rate”), number of fertile single crosses (“n_fertile_crosses”), rate of fertile crosses per survivors (“fertility_rate”), number of crosses that gave rise to transgenic F1 (“n_transgenic_G0”), and rate of transgenic founders per fertile crosses (“success rate”). Note that, for sequences, “left” means 5’ and “right” means 3’ in relation to the strand the gene is transcribed from.

**Supplementary File S2: Table in TXT format (tab-delimited) containing quantifications of transgenic marker removal**. Columns indicate tagged gene and terminus (“genotype”), a cross identifier (“cross”), number of screened flies (“N_screened”), number of screened flies without DsRed expression (“N_DsR-”), and the percentage of screened flies that lost DsRed expression (“ratio”).

**Supplementary File S3: Table in TXT format (tab-delimited) containing quantifications of tag exchange**. Columns indicate tagged gene (“genotype”), a cross identifier (“cross”), number of screened flies (“N_screened”), number of screened flies with gain of w and loss of DsRed expression (“N_w+_DsR-”), and the percentage of screened flies that gained w and lost DsRed expression (“ratio”).

## Data Availability

Plasmids will be deposited on Addgene and are currently available on request, as are generated fly lines.

Protocols and construct info will be available at http://www.TFTag.co.uk.

The bioinformatic pipeline for automated gRNA and homology arm design is explained in detail at https://github.com/emmacwatts/AutoTagsCRISPR/tree/main.

## Acknowledgements

We would like to thank Hilary Ashe, Simon Bullock, Cassandra Extavour, Eileen Furlong, Alistair McGregor, Alexandra Buffry, and Maike Kittelmann for discussions and helpful advice throughout the project. Synthetic DNA fragments were generated by Twist Bioscience and GenScript. Imaging was supported by the Oxford Brookes Centre for Bioimaging. Stocks obtained from the Bloomington *Drosophila* Stock Center (NIH P40OD018537) were used in this study. This project would not have been possible without the constant use of genomic sequences and gene annotations curated by FlyBase^74^.

## Author Contributions

Conceptualization: KH, SK; Data curation: EW, GB, ML, KH, SAH, SH, SK, SP, ZV; Formal analysis: EW, GB, ML, KH, SH, SK; Funding acquisition: KH, SK; Investigation: EW, GB, ML, KH, SAH, SH, SK, SP, ZV; Methodology: EW, GB, ML, KH, SH, SK, SP; Project administration: EW, GB, ML, KH, SAH, SH, SK, SP, ZV; Resources: EW, GB, ML, KH, SAH, SH, SK, SP, ZV; Software: EW, GB, ML; Supervision: KH, SK; Validation: EW, GB, ML, KH, SAH, SH, SK, SP, ZV; Visualization: EW, GB, ML, KH, SH, SK; Writing – original draft: EW, GB, ML, KH, SH, SK; Writing – review & editing: EW, GB, ML, KH, SAH, SH, SK, SP, ZV

## References

1. Gregor, T., Wieschaus, E. F., McGregor, A. P., Bialek, W. & Tank, D. W. Stability and nuclear dynamics of the bicoid morphogen gradient. Cell 130, 141–152 (2007).

2. Mir, M. et al. Dynamic multifactor hubs interact transiently with sites of active transcription in Drosophila embryos. eLife 7, e40497 (2018).

3. Trylinski, M., Mazouni, K. & Schweisguth, F. Intra-lineage Fate Decisions Involve Activation of Notch Receptors Basal to the Midbody in Drosophila Sensory Organ Precursor Cells. Curr Biol 27, 2239-2247.e3 (2017).

4. Bellen, H. J. et al. The Drosophila Gene Disruption Project: Progress Using Transposons With Distinctive Site Specificities. Genetics 188, 731–743 (2011).

5. Bischof, J. et al. A versatile platform for creating a comprehensive UAS-ORFeome library in Drosophila. Development 140, 2434–2442 (2013).

6. Kanca, O. et al. An efficient CRISPR-based strategy to insert small and large fragments of DNA using short homology arms. eLife 8, e51539 (2019).

7. Kanca, O. et al. An expanded toolkit for Drosophila gene tagging using synthesized homology donor constructs for CRISPR-mediated homologous recombination. Elife 11, e76077 (2022).

8. Kelso, R. J. et al. Flytrap, a database documenting a GFP protein-trap insertion screen in Drosophila melanogaster. Nucleic Acids Res 32, D418–420 (2004).

9. Kudron, M. M. et al. The ModERN Resource: Genome-Wide Binding Profiles for Hundreds of Drosophila and Caenorhabditis elegans Transcription Factors. Genetics 208, 937–949 (2018).

10. Lee, P.-T. et al. A gene-specific T2A-GAL4 library for Drosophila. eLife 7, e35574 (2018).

11. Lowe, N. et al. Analysis of the expression patterns, subcellular localisations and interaction partners of Drosophila proteins using a pigP protein trap library. Development 141, 3994–4005 (2014).

12. Nagarkar-Jaiswal, S. et al. A library of MiMICs allows tagging of genes and reversible, spatial and temporal knockdown of proteins in Drosophila. eLife 4, e05338 (2015).

13. Sarov, M. et al. A genome-wide resource for the analysis of protein localisation in Drosophila. eLife 5, e12068 (2016).

14. Venken, K. J. T. et al. MiMIC: a highly versatile transposon insertion resource for engineering Drosophila melanogaster genes. Nat Methods 8, 737–743 (2011).

15. Castells-Nobau, A. et al. Conserved regulation of neurodevelopmental processes and behavior by FoxP in Drosophila. PLoS One 14, e0211652 (2019).

16. Halder, G., Callaerts, P. & Gehring, W. J. Induction of ectopic eyes by targeted expression of the eyeless gene in Drosophila. Science 267, 1788–1792 (1995).

17. Minakuchi, C., Zhou, X. & Riddiford, L. M. Krüppel homolog 1 (Kr-h1) mediates juvenile hormone action during metamorphosis of Drosophila melanogaster. Mech Dev 125, 91– 105 (2008).

18. Scott, J. A., Williams, D. W. & Truman, J. W. The BTB/POZ zinc finger protein Broad-Z3 promotes dendritic outgrowth during metamorphic remodeling of the peripheral stretch receptor dbd. Neural Dev 6, 39 (2011).

19. Vavouri, T., Semple, J. I., Garcia-Verdugo, R. & Lehner, B. Intrinsic protein disorder and interaction promiscuity are widely associated with dosage sensitivity. Cell 138, 198–208 (2009).

20. Venken, K. J. T. et al. Versatile P[acman] BAC libraries for transgenesis studies in Drosophila melanogaster. Nat Methods 6, 431–434 (2009).

21. Diao, F. & White, B. H. A novel approach for directing transgene expression in Drosophila: T2A-Gal4 in-frame fusion. Genetics 190, 1139–1144 (2012).

22. Bateman, J. R., Lee, A. M. & Wu, C. Site-Specific Transformation of Drosophila via DC31 Integrase-Mediated Cassette Exchange. Genetics 173, 769–777 (2006).

23. Chen, Y.-C. D. et al. Using single-cell RNA sequencing to generate predictive cell-type-specific split-GAL4 reagents throughout development. Proc Natl Acad Sci U S A 120, e2307451120.

24. Forbes Beadle, L., Sutcliffe, C. & Ashe, H. L. A simple MiMIC-based approach for tagging endogenous genes to visualise live transcription in Drosophila. Development 151, dev204294 (2024).

25. Li, S. A., Li, H. G., Shoji, N., Desplan, C. & Chen, Y.-C. D. Protocol for replacing coding intronic MiMIC and CRIMIC lines with T2A-split-GAL4 lines in Drosophila using genetic crosses. STAR Protoc 4, 102706 (2023).

26. Li, L. et al. dFlpTag, a RMCE-based tool for simultaneous endogenous protein tagging and cell labeling in Drosophila. Sci Rep 15, 45542 (2025).

27. McKinney, H. M., Sherer, L. M., Williams, J. L., Certel, S. J. & Stowers, R. S. Characterization of Drosophila octopamine receptor neuronal expression using MiMIC-converted Gal4 lines. J Comp Neurol 528, 2174–2194 (2020).

28. Li-Kroeger, D. et al. An expanded toolkit for gene tagging based on MiMIC and scarless CRISPR tagging in Drosophila. Elife 7, e38709 (2018).

29. Bier, E., Harrison, M. M., O’Connor-Giles, K. M. & Wildonger, J. Advances in Engineering the Fly Genome with the CRISPR-Cas System. Genetics 208, 1–18 (2018).

30. Bruckner, J. J. et al. Fife organizes synaptic vesicles and calcium channels for high-probability neurotransmitter release. Journal of Cell Biology 216, 231–246 (2017).

31. Voutev, R. & Mann, R. S. Robust ΦC31-Mediated Genome Engineering in Drosophila melanogaster Using Minimal attP/attB Phage Sites. G3 Genes|Genomes|Genetics 8, 1399–1402 (2018).

32. Waldo, G. S., Standish, B. M., Berendzen, J. & Terwilliger, T. C. Rapid protein-folding assay using green fluorescent protein. Nat Biotechnol 17, 691–695 (1999).

33. Venken, K. J. T. et al. Genome engineering: Drosophila melanogaster and beyond. WIREs Developmental Biology 5, 233–267 (2016).

34. Port, F., Chen, H.-M., Lee, T. & Bullock, S. L. Optimized CRISPR/Cas tools for efficient germline and somatic genome engineering in Drosophila. Proc Natl Acad Sci U S A 111, E2967–2976 (2014).

35. Irion, U., Krauss, J. & Nüsslein-Volhard, C. Precise and efficient genome editing in zebrafish using the CRISPR/Cas9 system. Development 141, 4827–4830 (2014).

36. Zhang, J.-P. et al. Efficient precise knockin with a double cut HDR donor after CRISPR/Cas9-mediated double-stranded DNA cleavage. Genome Biol 18, 35 (2017).

37. Ren, X. et al. Optimized gene editing technology for Drosophila melanogaster using germ line-specific Cas9. Proc. Natl. Acad. Sci. U.S.A. 110, 19012–19017 (2013).

38. Thomsen, S., Azzam, G., Kaschula, R., Williams, L. S. & Alonso, C. R. Developmental RNA processing of 3′UTRs in Hox mRNAs as a context-dependent mechanism modulating visibility to microRNAs. Development 137, 2951–2960 (2010).

39. Huang, J., Zhou, W., Dong, W. & Hong, Y. Targeted engineering of the Drosophila genome. Fly (Austin) 3, 274–277 (2009).

40. Chou, T. & Perrimon, N. The Autosomal FLP-DFS Technique for Generating Germline Mosaics in Drosophila melanogaster. Genetics 144, 1673–1679 (1996).

41. Gohl, D. M. et al. A versatile in vivo system for directed dissection of gene expression patterns. Nat Methods 8, 231–237 (2011).

42. Bischof, J., Maeda, R. K., Hediger, M., Karch, F. & Basler, K. An optimized transgenesis system for Drosophila using germ-line-specific φC31 integrases. Proc. Natl. Acad. Sci. U.S.A. 104, 3312–3317 (2007).

43. Abu-Shaar, M. & Mann, R. S. Generation of multiple antagonistic domains along the proximodistal axis during Drosophila leg development. Development 125, 3821–3830 (1998).

44. Agrawal, P., Habib, F., Yelagandula, R. & Shashidhara, L. S. Genome-level identification of targets of Hox protein Ultrabithorax in Drosophila: novel mechanisms for target selection. Sci Rep 1, 205 (2011).

45. Delker, R. K., Ranade, V., Loker, R., Voutev, R. & Mann, R. S. Low affinity binding sites in an activating CRM mediate negative autoregulation of the Drosophila Hox gene Ultrabithorax. PLoS Genet 15, e1008444 (2019).

46. Garaulet, D. L. et al. Homeotic Function of Drosophila Bithorax-Complex miRNAs Mediates Fertility by Restricting Multiple Hox Genes and TALE Cofactors in the CNS. Developmental Cell 29, 635–648 (2014).

47. Gratz, S. J. et al. Highly Specific and Efficient CRISPR/Cas9-Catalyzed Homology-Directed Repair in Drosophila. Genetics 196, 961–971 (2014).

48. Ren, X. et al. Enhanced specificity and efficiency of the CRISPR/Cas9 system with optimized sgRNA parameters in Drosophila. Cell Rep 9, 1151–1162 (2014).

49. Port, F., Muschalik, N. & Bullock, S. L. Systematic Evaluation of Drosophila CRISPR Tools Reveals Safe and Robust Alternatives to Autonomous Gene Drives in Basic Research. G3 Genes|Genomes|Genetics 5, 1493–1502 (2015).

50. Skryabin, B. V. et al. Pervasive head-to-tail insertions of DNA templates mask desired CRISPR-Cas9-mediated genome editing events. Sci Adv 6, eaax2941 (2020).

51. Suchy, F. P. et al. Genome engineering with Cas9 and AAV repair templates generates frequent concatemeric insertions of viral vectors. Nat Biotechnol 43, 204–213 (2025).

52. Diao, F. et al. Plug-and-Play Genetic Access to Drosophila Cell Types using Exchangeable Exon Cassettes. Cell Reports 10, 1410–1421 (2015).

53. Nagarkar-Jaiswal, S. et al. A genetic toolkit for tagging intronic MiMIC containing genes. eLife 4, e08469 (2015).

54. Zirin, J. et al. Expanding the Drosophila toolkit for dual control of gene expression. eLife 12, RP94073 (2024).

55. Acevedo, J. M., Hoermann, B., Schlimbach, T. & Teleman, A. A. Changes in global translation elongation or initiation rates shape the proteome via the Kozak sequence. Sci Rep 8, 4018 (2018).

56. Aguilar, G. et al. Seamless knockins in Drosophila via CRISPR-triggered single-strand annealing. Developmental Cell 59, 2672-2686.e5 (2024).

57. Fleming, J. et al. AlphaFold Protein Structure Database and 3D-Beacons: New Data and Capabilities. Journal of Molecular Biology 437, 168967 (2025).

58. Jumper, J. et al. Highly accurate protein structure prediction with AlphaFold. Nature 596, 583–589 (2021).

59. Kim, A.-R. et al. FlyPredictome: A structural atlas of predicted protein-protein interactions in Drosophila. Preprint at 10.64898/2026.04.14.718529 (2026).

60. Caussinus, E., Kanca, O. & Affolter, M. Fluorescent fusion protein knockout mediated by anti-GFP nanobody. Nat Struct Mol Biol 19, 117–121 (2011).

61. Irizarry, J., McGehee, J., Kim, G., Stein, D. & Stathopoulos, A. Twist-dependent ratchet functioning downstream from Dorsal revealed using a light-inducible degron. Genes Dev 34, 965–972 (2020).

62. Trost, M., Blattner, A. C. & Lehner, C. F. Regulated protein depletion by the auxin-inducible degradation system in Drosophila melanogaster. Fly (Austin) 10, 35–46 (2016).

63. Götzke, H. et al. The ALFA-tag is a highly versatile tool for nanobody-based bioscience applications. Nat Commun 10, 4403 (2019).

64. Zhang, B., Zhang, Y. & Liu, J.-L. Highly effective proximate labeling in Drosophila. G3 (Bethesda) 11, jkab077 (2021).

65. Bothma, J. P., Norstad, M. R., Alamos, S. & Garcia, H. G. LlamaTags: A Versatile Tool to Image Transcription Factor Dynamics in Live Embryos. Cell 173, 1810-1822.e16 (2018).

66. Gilles, A. F., Schinko, J. B. & Averof, M. Efficient CRISPR-mediated gene targeting and transgene replacement in the beetle Tribolium castaneum. Development 142, 2832– 2839 (2015).

67. Koressaar, T. & Remm, M. Enhancements and modifications of primer design program Primer3. Bioinformatics 23, 1289–1291 (2007).

68. Untergasser, A. et al. Primer3—new capabilities and interfaces. Nucleic Acids Research 40, e115–e115 (2012).

69. Matinyan, N. et al. Multiplexed drug-based selection and counterselection genetic manipulations in Drosophila. Cell Rep 36, 109700 (2021).

70. Ni, J.-Q. et al. A genome-scale shRNA resource for transgenic RNAi in Drosophila. Nat Methods 8, 405–407 (2011).

71. Blair, S. S. Fixation of Imaginal Discs in Drosophila. Cold Spring Harb Protoc 2007, pdb.prot4795 (2007).

72. Spratford, C. M. & Kumar, J. P. Dissection and immunostaining of imaginal discs from Drosophila melanogaster. J Vis Exp 51792 (2014) doi:10.3791/51792.

73. Schindelin, J. et al. Fiji: an open-source platform for biological-image analysis. Nat Methods 9, 676–682 (2012).

74. Öztürk-Çolak, A. et al. FlyBase: updates to the Drosophila genes and genomes data-base. GENETICS 227, iyad211 (2024).

