## Supplementary tables and figures for "ExTaSy: A swappable CRISPR platform for endogenous tagging in *Drosophila melanogaster*"

### Supplementary materials

**Table S1: *Drosophila* lines.** For each line, the genotype, source and, if applicable, stock number, as well as notes on their use are given. Genotypes are indicated as per https://flybase.org/ nomenclature. BDSC = Bloomington *Drosophila* Stock Center.

| **Genotype** | **Source/stock number (if applicable)** | **Use/notes** |
|---|---|---|
| y[1] w[1118] | BDSC_6598 | Outcrossing of injected flies |
| svb[PL107]/FM0 | Francois Payre, Toulouse | Balancing of insertions |
| y[1] w[*]; wg[Sp-1]/CyO; Dr[1]/TM3, Sb[1] | BDSC_59967 | Balancing of insertions |
| y[1] w[*]; TI{RFP[DsRed.3xP3.cUa]=TI}Crk[dsRed]/TI{GMR-HMS04515}Gat[eya] | BDSC_90850 | Balancing of insertions |
| y[1] sc[*] v[1] sev[21]; P{y[+t7.7] v[+t1.8]=nanos-Cas9.R}attP40 | BDSC_78781 | CRISPR/Cas9 for loci on *chr3*; Ren et al., 2013 |
| y[1] sc[*] v[1] sev[21]; P{y[+t7.7] v[+t1.8]=nanos-Cas9.R}attP2 | BDSC_78782 | CRISPR/Cas9 for loci on *chr2* and *chr4*; Ren et al., 2013 |
| y[1] w[*]; P{y[+t7.7] v[+t1.8]=nanos-Cas9.R}attP2 | This study | BDSC_59967 crossed to BDSC_78782; CRISPR/Cas9 for loci on *chrX* |
| P{ry[+t7.2]=hsFLP}12, y[1] w[*] M{vas-int.B}ZH-2A; Pri[1]/TM3, Sb[1] | BDSC_33216 | Expression of Flp and ΦC31 integrase for RMCE; constructs recombined on *chrX*; Chou & Perrimon, 1996; Gohl *et al.*, 2011 |
| y[1] w[*]; P{GFP[3xP3.cLa]=hsFLP, vas.int}zh51; Dr[1]/TM3, Sb[1] | This study | Expression of Flp and ΦC31 integrase for RMCE; constructs inserted on *chr2* |
| w[1118]; CyO, P{Tub-PBac\T}2/wg[Sp-1]; l(3)*[*]/TM6B, Tb[1] | BDSC_8285 | Expression of pBac transposase for transgenic marker removal |
| y[1] w[*]; P{w[+mC]=sfGFP.swap}Zh30, P{w[+mC]=sfGFP.swap}Zh51, P{w[+mC]=sfGFP.swap}Zh58; Dr[1]/TM3, Sb[1] | This study | Swap line for exchange of 3XHA tag at the C-terminus for sfGFP; construct inserted on *chr2* |

| | | |
|---|---|---|
| y[1] w[*]; wg[Sp-1]/CyO; P{w[+mC]=sfGFP.swap}Zh64, P{w[+mC]=sfGFP.swap}Zh86, P{w[+mC]=sfGFP.swap}Zh96 | This study | Swap line for exchange of 3XHA tag at the C-terminus for sfGFP; construct inserted on *chr3* |
| y[1] w[*]; pExTaSy-EC{RFP[3xP3.cLa]=3xHA}Ubx/TM3, Sb[1] | This study; TFTag_04B10 | 3XHA tag at C-terminus of *Ubx*; tagging all protein isoforms |
| y[1] w[*]; pSwap-C{w[+mC]=sfGFP}Ubx[12.3-1]/TM3, Sb[1] | This study | Line TFTag_04B10 where original 3XHA tag was exchanged for sfGFP |
| y[1] w[*]; pExTaSy-EN{RFP[3xP3.cLa]=3xHA}Ubx[3.1]/TM3, Sb[1] | This study; TFTag_04B11 | 3XHA tag at N-terminus of *Ubx*; tagging all protein isoforms |
| y[1] w[*]; pExTaSy-EN{3xHA}Ubx[3.1-11.1] | This study | Line TFTag_04B11 where DsRed transgenic marker has been removed |
| y[1] w[*]; pExTaSy-EC{RFP[3xP3.cLa]=3xHA}hth[3.3] | This study; TFTag_05B01 | 3XHA tag at C-terminus of *hth*; tagging protein isoform E (FBpp0099878) |
| y[1] w[*] pExTaSy-ECRFP[3xP3.cLa]=3xHA}exd[1a.4]; wg[Sp-1]/CyO | This study; TFTag_05B09 | 3XHA tag at C-terminus of *exd*; tagging all protein isoforms |

**Table S2: Synthetic DNA fragments**. We give the sequence for each fragment and indicate which plasmid it was used for. We also indicate which components of the final plasmid are on the fragment.

| plasmid | sequence | components |
|---|---|---|
| *pExTaSy-EC* | TTGAAAAAGTGGCACCGAGTCGGTGCTAACAAAGCACCAGT GGTCTAGTGGTAGAATAGTACCCTGCCACGGTACAGACCCG GGTTCGATTCCCGGCTGGTGCAGGAAGAGCCGTCGCTCTTC CGGATCCGCAGGAAGTGCCGCTGGCAGCGGTGAATTCGGT GCGGGTGCCAGGGCGTGCCCCCGGGCTCCCCGGGCGCG TACTCCATACCCGTATGACGTTCCTGACTACGCGGGGTATCC CTATGACGTGCCGGACTATGCGGGGTCTTACCCCTACGACG TGCCAGACTACGCTTAATTAACCCTAGAAAGATAATCATATTGT GACGTACGTTAAAGATAATCATGCGTAAAATTGACGCATGTGT TTTATCGGTCTGTATATCGAGGTTTAT | Rice tRNA:Gly, SapI fragment, linker, *attB*$^{CC}$ site, 3XHA tag, 3’ end of right (3’) pBac inverted repeat |
| *pExTaSy-EC* | TTAGAAAGAGAGAGCAATATTTCAAGAATGGTGCCCCAACTG GGGTAACCTTTGAGTTCTCTCAGTTGGGGGCATGCGTCAATTT TACGCAGACTATCTTTCTAGGGTTAAGGAAGAGCCGTCGCTC TTCCTCATGTTCTGATCGGAACGACCGATCCTCGCCTCAGTT AGCGTACATCCTACCTATACTGCTTCCTATACGACATCACCG ATG | *attP* site, 3’ end of left (5’) pBac inverted repeat, SapI fragment, |

| | | |
|---|---|---|
| | | spacer, sgRNA-synth binding site |
| *pExTaSy-EN* | TTGAAAAAGTGGCACCGAGTCGGTGCTAACAAAGCACCAGT<br>GGTCTAGTGGTAGAATAGTACCCTGCCACGGTACAGACCCG<br>GGTTCGATTCCCGGCTGGTGCAGGAAGAGCCGTCGCTCTTC<br>CTTAACCCTAGAAAGATAATCATATTGTGACGTACGTTAAAGAT<br>AATCATGCGTAAAATTGACGCATGTGCGGGTGCCAGGGCGT<br>GCCCTTGGGCTCCCCGGGCGCGTACTCCGGGTGTTTTATCG<br>GTCTGTATATCGAGGTTTAT | Rice tRNA:Gly, SapI fragment, 3' end of right (3') pBac inverted repeat, *attB* site |
| *pExTaSy-EN* | ATTTTACGCAGACTATCTTTCTAGGGTTAAATGTACCCGTATGA<br>CGTTCCTGACTACGCGGGGTATCCCTATGACGTGCCGGACT<br>ATGCGGGGTCTTACCCCTACGACGTGCCAGACTACGCTGG<br>GTGCCCCAACTGGGGTAACCTCCGAGTTCTCTCAGTTGGGG<br>GAGGATCCGCAGGAAGTGCCGCTGGCAGCGGTGAATTCGG<br>AAGAGCCGTCGCTCTTCCTCATGTTCTGATCGGAACGACCG<br>ATCCTCGCCTCAGTTAGCGTACATCCTACCTATACTGCTTCCT<br>ATACGACATCACCGATG | 3XHA tag, *attP*$^{CC}$ site, linker, SapI fragment, spacer, sgRNA-synth binding site |
| *pExTaSy-EC and pExTaSy-EN* | CCATCGGTGATGTCGTATAAGAGACGTATATTTTTTGCTCACC<br>TGTGATTGCTCCTACTCAAATACAAAAACATCAAATTTTCTGTC<br>AATAAAGCATATTTATTTATATTTATTTTACAGGAAAGAATTCCTT<br>TTAAAGTGTATTTTAACCTATAATGAAAAACGATTAAAAAAAATA<br>CATAAAATAATTCGAAAATTTTTGAATAGCCCAGGTTGATAAAA<br>ATTCATTTCATACGTTTTATAACTTATGCCCCTAAGTATTTTTTGA<br>CCATAGTGTTTCAATTCTACATTAATTTTACAGAGTAGAATGAAA<br>CGCCACCTACTCAGCCAAGAGGCGAAAAGGTTAGCTCGCC<br>AAGCAGAGAGGGCGCCAGTGCTCACTACTTTTTATAATTCTCA<br>ACTTCTTTTTCCAGACTCAGTTCGTATATATAGACCTATTTTCAA<br>TTTAACGTCGGGGCTTTGAGTGTGTGTAGACATCAAGCATCG<br>GTGGTTCAGTGGTAGAATGCTCGCCTGCCACGCGGGCGGC<br>CCGGGTTCGATTCCCGGCCGATGCAGGTAGGATGTACGCTA<br>ACTGGTTTTAGAGCTAGAAATAGCAAGTTAAAATAAGGCTAGT<br>CCGTTATCAACTTGAAAAAGTGGCACCGAGTCGGTGCTAAC | *Drosophila U6* promoter, tRNA:Gly-GCC-1-1, sgRNA-synth sequence, gRNA scaffold |
| *pFlpPhi* | TCAGCCTAGGGTAGGATGTACGCTAACTGAGGTTTAAACCGC<br>GGGACGTCGAATCCCAAAACAAACTGGTTATTGTGGTAGGTC<br>ATTTGTTTGGCAGAAAGAAAACTCGAGAAATTTCTCTGGCCGT<br>TATTCGTTATTGTCTCTTTTCTTTTTGGGTCTCTCCCTCTCTGCA<br>CTAATGCTCTCTCACTCTGTCACACAGTAAACGACATACTGCT<br>CTCGTTGGTTCGAGAGAGCGCGCCTCGAATGTTCGCGAAAA<br>GAGCGCCGGAGTATAAATAGAGGCGCTTCGTCTACGGAGCG<br>ACAATTCAATTCAAACAAGCAAAGTGAACACGTCGCTAAGCG<br>AAAGCTAAGCAAATAAACAAGCGCAGCTGAACAAGCTAAACA<br>ATCTGCAGTAAAGTGCAAGTTAAAGTGAATCAATTAAAAGTAA<br>CCAGCAACCAAGTAAATCAACTGCAACTACTGAAATCTGCCA<br>AGAAGTAATTATTGAATACAAGAAGAGAACTCTGAATACTTTCA | *Drosophila* codon optimized *flippase* CDS, *hsp70Bb* promoter element, *AcNPV p10* 3'UTR, sgRNA-synth1 |

| | | |
|---|---|---|
| | ACAAGTTACCGAGAAAGAAGAACTCACACACAATGCCCCAG TTCGACATACTGTGTAAAACGCCCCCAAAAGTCCTCGTCCGT CAGTTCGTCGAAAGGTTCGAACGCCCGTCGGGCGAAAAAAT AGCACTGTGCGCCGCAGAACTGACCTATCTCTGCTGGATGA TCACGCATAATGGTACAGCAATCAAGCGGGCGACCTTTATGT CCTACAACACCATAATATCCAACTCCTTGAGCTTTGATATCGT AAACAAATCGCTCCAGTTTAAATACAAAACGCAGAAAGCGAC GATTTTGGAAGCTAGCCTCAAAAAACTCATCCCCGCGTGGGA ATTCACAATAATCCCTTACTACGGCCAAAAGCACCAGTCGGA CATAACAGACATAGTTAGTAGTCTCCAGTTGCAGTTCGAGTCG AGTGAGGAAGCGGACAAAGGCAATTCGCACAGCAAGAAGAT GCTCAAAGCCCTCCTGAGCGAAGGCGAATCCATCTGGGAG ATCACGGAGAAGATTCTGAATAGCTTCGAATATACAAGTCGGT TTACCAAGACAAAGACTTTGTACCAATTTTTGTTCCTCGCAACT TTTATCAATTGTGGTAGGTTCTCCGATATAAAGAACGTCGATC CTAAGAGCTTCAAGCTCGTACAGAATAAATACCTCGGCGTAA TTATTCAGTGCCTCGTCACAGAGACTAAAACCTCGGTTAGTC GTCATATATACTTCTTTTCGGCCCGTGGACGAATCGATCCATT GGTATATCTCGACGAATTTTTGCGTAATTCCGAGCCAGTTTTG AAGCGTGTAAACAGGACTGGTAATAGTTCGAGCAACAAACAG GAATATCAACTGCTCAAGGACAACCTGGTTAGGAGCTATAATA AGGCACTCAAGAAGAACGCACCGTATAGTATCTTTGCGATAA AAAACGGTCCAAAGAGCCACATCGGACGGCATTTGATGACC AGCTTCCTGAGTATGAAGGGTCTCACAGAACTCACCAACGTC GTGGGTAACTGGTCCGATAAGAGGGCATCCGCCGTTGCCC GCACGACCTATACACATCAAATCACAGCGATCCCTGATCACT ACTTCGCCCTCGTATCGCGCTATTATGCGTACGATCCAATAT CCAAGGAGATGATCGCCCTCAAGGACGAAACGAACCCTATC GAGGAGTGGCAGCACATAGAGCAGTTGAAAGGCAGTGCCG AGGGTTCGATCCGATACCCTGCATGGAATGGAATTATCTCGC AAGAAGTGTTGGATTATCTCTCCTCCTATATAAATAGGCGAATA TAACTAGAATGAATCGTTTTTAAAATAACAAATCAATTGTTTTATA ATATTCGTACGATTCTTTGATTATGTAATAAAATGTGATCATTAG GAAGATTACGAAAAATATAAAAAATATGAGTTCTGTGTGTATAA CAAATGCTGTAAACGCCACAATTGTGTTTGTTGCAAATAAACC CATGATTATTTGATTAAAATTGTTGTTTTCTTTGTTCATAGACAAT AGTGTGTTTTGCCTAAACGTGTACTGCATAAACTCCATGCGAG TGTATAGCGAGCTAGTGGCTAACGCTTGCCCCACCAAAGTA GATTCGTCAAAATCCTCAATTTCATCACCCTCCTCCAAGTTTA ACATTTGGCCGTCGGAATTAACTTCTAAAGATGCCACATAATC TAATAAATGAAATAGAGATTCAAACGTGGCGTCATCGTCCGTT TCGACCATTTCCGAAAAGAACTCGGGCATAAACTCTATGATTT CTCTGGACGTGGTGTTGTCGAAACTCTCAAAGTACGCAGTCA GGAACGTGCGCGACATGTCGTCGGGAAACTCGCGCGGAAA CATGTTGTTGTAACCGAACGGGTCCCATAGCGCCAAAACCA AATCTGCCAGCGTCAATAGAATGAGCACGATGCCGACAATG GAGCTGGCTTGGATAGCGATTCGAGTTAACGGCCGGGCCCT GAC | recognition sequence |
| *pFlpPhi* | GTAATACGACTCACTATAGGGCGAATTGGGTACGGTAGGATG TACGCTAACTGAGGGTGGATTGGATTGTCATTTTTACGTAACTA | *vasa* promoter |

| | | |
|---|---|---|
| | GTGCGGCCGCGTAGTTTGGGTTTCAGAAGTCAACAGTGTTAC<br>CGTATTTTAATGTACATCTGGCTTTTAAGCGAGTACACACACAT<br>AGCATTGCGATGTTTTTTAAACTTGCACTGCTTGCAAGCAGGG<br>CTCCACTGTATTTCAATGAAAGAACAGCGTGTTGGGCGTTTTT<br>GAATTATACTAATTACCTTCAATAACAATTCCCTATATATCACTT<br>AGTTTTAATAAATAAATCGTTTGAGTGTGGTCTAGAGATGATGT<br>ATTATGATGAAGTGCGGTTGTTGGGGACAAGCGGGATACGTC<br>TGCGGGTCAAATCGGCGGTGGGACATAGTGAGAGGGCAAC<br>GGCTCCAAAACCAAGCGTTGCGAACAATCACCAACGCGCC<br>CTGGTACGCGAGAAATGTTGATATCGCACATGACCTCAGCAT<br>CCACCTTCAGTTTGAGAACAAATTAAATCAGGCACATACCAG<br>ACAACCAGGAGAACACTGAGGAGGCGTCATCAAAGATATCTT<br>TGGACACGTGGCATAAACAAGCCAAACTATTATCTAATCAATA<br>TTTTTATACAAATTTTGTATGTTTCCATGTTATCTTCGTTGCTCCT<br>GTTAGTTACTACGTTTCATAGCCCTTAAATTGGTTGCTTAACGT<br>AAAATAAAATTATATTGTATAAAAATAAAGAGGAATTCACCTATA<br>ACAAACGAAAAAGTAAATAAATAAGAACTTCCCCATGTTAAAA<br>ATGCCACCACCATCTTTACATATAAGTATGTACATATGAATGAA<br>TCACTTAGGTTGCTTGAATATTACAATTTCATTAATAGCTAAATC<br>ACATTTGATGTGTTAGTGGAAAACGGCTATATATATATAAATTATC<br>GAAATTGTGAATATCGAATTGCGATAGCACAATGGGAAATTCC<br>ACCACTAGATTTTTGGTACTTTTAACAGATCCTTTTCGGTTTTGC<br>GTTGCGCGAAGTGATCTGAACTTATCAAAGTTTGTAAGGTAAT<br>ACATAAAGTGAAAAAGAATTAATTTGCTCTTGAAAGGCAGGCC<br>AAATTAAAAAAAAAATATCAATATGGACACGTATGCCGGTGCT<br>TACGACCGTC | |
| *pFlpPhi* | ATGGACACGTATGCCGGTGCTTACGACCGTCAGAGTAGGGA<br>ACGGGAAAACAGCAGCGCCGCCAGTCCCGCGACGCAACG<br>GAGCGCGAACGAGGACAAAGCTGCTGATTTGCAGCGAGAG<br>GTGGAACGAGATGGTGGTCGTTTCAGGTTCGTCGGACATTTT<br>AGTGAAGCCCCAGGCACGAGCGCGTTCGGTACTGCCGAGC<br>GTCCAGAGTTCGAACGAATACTGAACGAGTGTAGGGCGGGA<br>CGTCTGAACATGATAATCGTATACGACGTGAGTCGCTTCAGC<br>CGATTGAAAGTTATGGACGCAATACCGATAGTCAGTGAACTCT<br>TGGCCCTGGGTGTCACTATAGTCAGCACTCAAGAGGGAGTAT<br>TTCGTCAAGGTAACGTAATGGACCTCATCCACCTCATTATGC<br>GGTTGGACGCCAGCCATAAAGAGAGTAGCCTCAAAAGCGC<br>CAAGATACTCGACACGAAGAATCTGCAGCGGGAACTGGGC<br>GGTTACGTCGGTGGAAAGGCTCCATATGGCTTTGAATTGGTG<br>TCCGAGACAAAGGAAATTACGCGTAATGGCCGAATGGTCAAT<br>GTTGTGATAAACAAGCTCGCACATTCGACGACCCCGCTCAC<br>AGGTCCATTCGAATTTGAGCCGGACGTTATTAGGTGGTGGTG<br>GCGGGAGATCAAGACACATAAACACCTGCCGTTTAAACCCG<br>GAAGTCAGGCCGCTATACATCCAGGCAGTATAACAGGACTG<br>TGTAAGCGTATGGACGCCGATGCTGTCCCCACTCGAGGTGA<br>AACGATTGGTAAGAAGACTGCAAGTTCGGCTTGGGATCCTGC<br>AACAGTGATGCGAATTCTCCGCGACCCTCGGATAGCGGGAT<br>TCGCGGCCGAGGTAATATACAAGAAAAAGCCAGATGGAACC<br>CCTACTACTAAGATTGAGGGATATCGCATTCAGCGTGATCCG<br>ATTACACTGCGCCCTGTCGAATTGGATTGCGGACCAATTATA | *Drosophila* codon-optimized *PhiC31* |

| | | |
|---|---|---|
| | GAGCCCGCAGAGTGGTATGAGCTCCAAGCGTGGCTGGATG GACGTGGAAGGGGAAAAGGTCTCAGTCGCGGACAGGCCAT TTTGTCGGCGATGGACAAACTCTATTGCGAATGTGGAGCTGT CATGACATCCAAACGAGGCGAGGAGAGTATTAAAGACTCGTA TCGCTGCCGTCGCCGGAAGGTCGTCGATCCCTCGGCGCCC GGCCAGCATGAAGGCACATGCAACGTGTCGATGGCAGCCTT GGACAAATTCGTTGCCGAGAGGATTTTCAACAAAATTAGGCAT GCCGAGGGAGATGAGGAAACTCTCGCCTTGTTGTGGGAGGC GGCACGACGATTCGGTAAGCTCACAGAGGCGCCGGAGAAG TCGGGCGAGAGGGCGAATTTGGTGGCAGAGCGGGCGGATG CTCTCAATGCTCTCGAGGAACTGTACGAAGACCGAGCAGCA GGTGCGTACGACGGTCCAGTTGGCCGGAAACATTTCCGCAA ACAACAGGCTGCATTGACATTGCGCCAACAAGGTGCCGAGG AGCGTTTGGCTGAGTTGGAAGCGGCTGAAGCTCCTAAGTTGC CCCTCGATCAATGGTTCCCGGAAGATGCCGATGCAGATCCG ACTGGTCCGAAGTCGTGGTGGGGTCGCGCTTCCGTCGATGA CAAGCGTGTATTTGTTGGCTTGTTCGTTGATAAAATAGTGGTCA CTAAGAGCACGACGGGTAGGGGACAGGGAACGCCTATAGA GAAACGGGCTTCGATCACGTGGGCCAAGCCGCCGACCGAC GACGATGAGGATGACGCCCAGGATGGCACCGAAGACGTAG CGGCCTAG | |
| *pFlpPhi* | CAGGATGGCACCGAAGACGTAGCGGCCTAGAATGTATGGAC ATAGATTTCAAATAATTAAATGTAATGCAGTAATTGATGTAATTA GTTAAATAAGTTAGATATTAATAACATATTAATTATATGTATTATAA CGCATATAATAATAAAATGCATATTTAGGATATGCAAGCCATTC GAATTTTCTATTTTAATTTCCTTTTACAAAGAAATGTATAACAAAA TATAATTTGAAAAAATGTTCTGGCTCTAATTCGATTTCTTTTAAGT ATTTTGTGAACTGCTTTTAATAACGAGCGGTTGCAAAACTTAAC AGAAACGTGCACTTTGATCCCAACTAATACCGTGTACTATCAC GTGTTTGGTTTTAAGTACTCATCCTTCGTGTTTCGTTGTTTGTTG TTTGCCTTGCTGTACACTTGGCTCTTGCGCTCTCTCGCTCTCC GTTGGAGCCGGCTTTTTGAATGCATGCCTCGCCCTGCTCGC GCCCAGTCCTCCTCCCCCGAAAAGCATGCCGACCAGCAAA TGTTGCCCTTTTGCGTTTGCTGTCTTTGCAGAAGCAAAATCAAT AACTGAGAAATCCACCACACTGCTGCTCTTCGTGTTACCGGT GTACCGGGCCGCAGCTTTTTTTGACCACTTAG | *vasa* 3’ sequences |
| *pFlpPhi_GFP* | AAATCCACCACACTGCTGCTCTTCGTGTTACCGGTGGATCTA ATTCAATTAGAGACTAATTCAATTAGAGCTAATTCAATTAGGAT CCAAGCTTATCGATTTCGAACCCTCGACCGCCGGAGTATAAA TAGAGGCGCTTCGTCTACGGAGCGACAATTCAATTCAAACAA GCAAAGTGAACACGTCGCTAAGCGAAAGCTAAGCAAATAAA CAAGCGCAGCTGAACAAGCTAAACAATCGGCTCGAAGCCG GTCGCCACCAGATCTAAAAGGTAGGTTCAACCACTGATGCCT AGGCACACCGAAACGACTAACCCTAATTCTTATCCTTTACTTC AGGCGGCCGCGGCTCGAGAATCAAAATGGGCAACAAATGC TGCAGCAAGCGACAGGATCAGGAACTGGCACTGGCCTATC CCACTGGGGGCTACAAGAAATCCGACTACACCTTTGGCCAG ACGCACATCAACAGCAGCGGCGGCGGCAACATGGGCGGC GTTCTTGGCCAGAAGCATAACAACGGTGGCTCGCTGGACTC | *3xP3-GFP* transgenic marker |

| | |
|---|---|
| | GCGCTACACGCCCGATCCCAATCATCGGGGTCCGTTGAAAA TCGGCGGAAAGGGCGGCGTTGACATCATCAGACCACGCGG ATCCATGTCCAAAGGTGAAGAACTGTTTACCGGAGTAGTCCC GATATTGGTTGAACTCGACGGCGATGTCAACGGTCATAAATTC AGTGTGTCCGGCGAGGGTGAGGGCGACGCCACATACGGTA AGCTGACGTTGAAGTTCATATGCACCACGGGCAAGCTGCCC GTGCCATGGCCGACGTTGGTCACGACGCTGACGTATGGTGT CCAGTGTTTCAGCCGTTACCCCGATCATATGAAGCAGCATGA CTTTTTCAAGTCGGCGATGCCGGAGGGATACGTTCAAGAGAG GACCATTTTCTTCAAGGATGACGGCAACTATAAGACGCGAGC GGAGGTGAAATTTGAAGGCGACACACTCGTTAACCGTATTGA GTTGAAGGGCATTGATTTTAAGGAGGATGGCAACATTCTGGG CCATAAGTTGGAATACAACTACAACTCGCATAATGTGTATATAA TGGCAGATAAGCAGAAAAACGGAATAAAGGTTAACTTCAAGA TTCGCCACAACATAGAGGACGGTTCCGTGCAACTTGCAGAT CATTACCAACAGAACACACCCATTGGAGATGGCCCAGTTCTC TTGCCAGACAATCACTACCTTTCCACACAGTCCGCGTTGAGC AAGGACCCCAATGAAAAGCGGGACCACATGGTGTTGCTGGA GTTTGTGACCGCAGCTGGTATTACACACGGCATGGATGAGCT CTACAAGTAATCTAGAGGATCTTTGTGAAGGAACCTTACTTCT GTGGTGTGACATAATTGGACAAACTACCTACAGAGATTTAAAG CTCTAAGGTAAATATAAAATTTTTAAGTGTATAATGTGTTAAACTA CTGATTCTAATTGTTTGTGTATTTTAGATTCCAACCTATGGAACT GATGAATGGGAGCAGTGGTGGAATGCCTTTAATGAGGAAAAC CTGTTTTGCTCAGAAGAAATGCCATCTAGTGATGATGAGGCTA CTGCTGACTCTCAACATTCTACTCCTCCAAAAAAGAAGAGAA AGGTAGAAGACCCCAAGGACTTTCCTTCAGAATTGCTAAGTTT TTTGAGTCATGCTGTGTTTAGTAATAGAACTCTTGCTTGCTTTG CTATTTACACCACAAAGGAAAAAGCTGCACTGCTATACAAGA AAATTATGGAAAAATATTTGATGTATAGTGCCTTGACTAGAGAT CATAATCAGCCATACCACATTTGTAGAGGTTTTACTTGCTTTAA AAAACCTCCCACACCTCCCCCTGAACCTGAAACATAAAATGA ATGCAATTGTTGTTGTTAACTTGTTTATTGCAGCTTATAATGGTT ACAAATAAAGCAATAGCATCACAAATTTCACAAATAAAGCATTT TTTTCACTGCATTCTAGTTGTGGTTTGTCCAAACTCATCAATGT ATCTTATCATGTCTGGAACTAGTTAGAGGGCCCGGCCGTTAA CTCGAATCGC |

**Table S3: PCR primers.** Primer name, sequence (5’-3’), and use are indicated.

| **Primer name** | **sequence** |
|---|---|
| pExTaSy-E_bb_F1 | TATACTGCTTCCTATACGACATCAC |
| pExTaSy-E_bb_R1 | TATACGTCTCTTATACGACATCACCG |
| pExTaSy-E_ins_F1 | TGTTTTATCGGTCTGTATATCGAGG |
| pExTaSy-E_ins_R1 | CATTCTTGAAATATTGCTCTCTCTTTC |
| N-term_DsRed-rev | TTAACCCTAGAAAGATAGTCTGC |

| | |
|---|---|
| GMR-W-hsp70-F1 | CAATTGTTGTTGTTAACTTGTTTATTGC |
| GMR-W-hsp70-R1 | ACTAGCACCTGACTGTCGAG |
| Acc65I-hsp70prom-F1 | TTAGGGTACCGCTAGAATCCCAAAACAAACTGG |
| NheI-TERM-R1 | GTTGGCTAGCCTCCTGACAGCGGAACAAAC |
| zh30A_HAL_F1 | TGGATTGGATTGTCATTTTGTATACCGTGAGTTCGTTTTCCATTTTAAGCTGTCG |
| zh30A_HAL_R1 | TTCGGAATAGGAACTTCGGACTAGTGTGGTTCTATTTGGGCATCGGGTGCTTG |
| zh30A_HAR_F1 | AGAAAGTATAGGAACTTCTTGGAATTCTGATGGTGGTGTCCTTTGAAATGTTG |
| zh30A_HAR_R1 | GGATGTACGCTAACTGAGGGCCCGGGCGGTGCAGTCAAGGCCAGTTGAAC |
| zh51D_HAL_F1 | TGGATTGGATTGTCATTTTGTATACCGTGAACTGGGACTCACGGTTCAAC |
| zh51D_HAL_R1 | TTCGGAATAGGAACTTCGGACTAGTGTGGTTGTGGGCAGACAGCGGATG |
| zh51D_HAR_F1 | AGAAAGTATAGGAACTTCTTGGAATTCTGACTGAGGCGACTCCAACGC |
| zh51D_HAR_R1 | GGATGTACGCTAACTGAGGGCCCGGGCGGTCTGTGTACATTCCCGCACC |
| zh58A_HAL_F1 | TGGATTGGATTGTCATTTTGTATACCGTGAGTTGGGCCGAGGAAATTGTG |
| zh58A_HAL_R1 | TTCGGAATAGGAACTTCGGACTAGTGTGGTATCCACAGTCACCCACCATAAAG |
| zh58A_HAR_F1 | AGAAAGTATAGGAACTTCTTGGAATTCTGAGTGTGGAATTCAATGGGATGTTC |
| zh58A_HAR_R1 | GGATGTACGCTAACTGAGGGCCCGGGCGGTGCAACGGGGCCTTAATTTGG |
| zh64A_HAL_F1 | TGGATTGGATTGTCATTTTGTATACCGTGATGCGACAGTTTCCGTTTTGC |
| zh64A_HAL_R1 | TTCGGAATAGGAACTTCGGACTAGTGTGGTGCGAGGGGCAAATTTTAAGTTTC |
| zh64A_HAR_F1 | AGAAAGTATAGGAACTTCTTGGAATTCTGATTCCTTTTTCACCGTGCAGGATATAG |

| | |
|---|---|
| zh64A_HAR_R1 | GGATGTACGCTAACTGAGGGCCCGGGCGGTGGTCGCAGCCTCAGCATC |
| zh86Fb_HAL_F1 | TGGATTGGATTGTCATTTTGTATACCGTGAGCTTATCTTTTGTCGCGAGCTG |
| zh86Fb_HAL_R1 | TTCGGAATAGGAACTTCGGACTAGTGTGGTTCAGGTGTTGTTGCTATTTGGAC |
| zh86Fb_HAR_F1 | AGAAAGTATAGGAACTTCTTGGAATTCTGAGCGCGGTTAGAAATTATTCAGG |
| zh86Fb_HAR_R1 | GGATGTACGCTAACTGAGGGCCCGGGCGGTACGGCAATTAACACCTTGTGC |
| zh96E_HAL_F1 | TGGATTGGATTGTCATTTTGTATACCGTGACGAGGCTCATTGGCTTTTCG |
| zh96E_HAL_R1 | TTCGGAATAGGAACTTCGGACTAGTGTGGTGCACGGCTCACGTTTGCC |
| zh96E_HAR_F1 | AGAAAGTATAGGAACTTCTTGGAATTCTGAAGCACGTTAATCGATTCAAATCG |
| zh96E_HAR_R1 | GGATGTACGCTAACTGAGGGCCCGGGCGGTGACTTTTGCCGGCTCACAAG |
| SwapSyC_sfGFP_F1 | GGTAACCTCCGAGTTCTCTCAGTTGGGGGAGTGTCCAAGGGCGAGGAG |
| SwapSyC_sfGFP_R1 | GTCACAATATGATTATCTTTCTAGGGTTAATTACTTGTACAGCTCATCCATGC |
| SwapSyN_sfGFP_F1 | ATTTTACGCAGACTATCTTTCTAGGGTTAAATGGTGTCCAAGGGCGAGGAG |
| SwapSyN_sfGFP_R1 | AGCCCGGGGGCACGCCCTGGCACCCGCACCCTTGTACAGCTCATCCATGCC |
| pExTaSy_L_Seq_F1 | GTGTAGACATCAAGCATCGG |
| pExTaSy_C_Seq_F1 | AAGCGGCGACTGAGATGTC |
| SpeI-gypsy_F1 | TGGGACTAGTTGGCCACGTAATAAGTGTGCG |
| NotI-gypsy_R1 | CTACGCGGCCGCGTTGTTGGTTGGCACACC |
| AatII-gypsy_F1 | ATTCGACGTCTGGCCACGTAATAAGTGTGCG |
| SacII-gypsy_R1 | TAGACCGCGGGTTGTTGGTTGGCACACC |
| FlpPhi_zh2A_HAL-F1 | ACTGAGGGTGGATTGGATTGTCATTTTTACCAAACCACACCACCACACAC |

| | |
|---|---|
| FlpPhi_zh2A_HAL-R1 | CGCACACTTATTACGTGGCCAACTAGTTACACACCCGGGTTTCACATAAATAC |
| FlpPhi_zh2A_HAR-F1 | TGTGGTGTGCCAACCAACAACCCGCGGTTTGTATTTCGTTTCCCGCCATCTTG |
| FlpPhi_zh2A_HAR-R1 | CTAGGGTAGGATGTACGCTAACTGAGGTTTGTACACGGTGAAGGAGGTGG |
| FlpPhi_zh51D_HAL-F1 | ACTGAGGGTGGATTGGATTGTCATTTTTACACTGGGACTCACGGTTCAAC |
| FlpPhi_zh51D_HAL-R1 | CGCACACTTATTACGTGGCCAACTAGTTACTGTGGGCAGACAGCGGATG |
| FlpPhi_zh51D_HAR-F1 | TGTGGTGTGCCAACCAACAACCCGCGGTTTCTGAGGCGACTCCAACGC |
| FlpPhi_zh51D_HAR-R1 | CTAGGGTAGGATGTACGCTAACTGAGGTTTTCTGTGTACATTCCCGCACC |

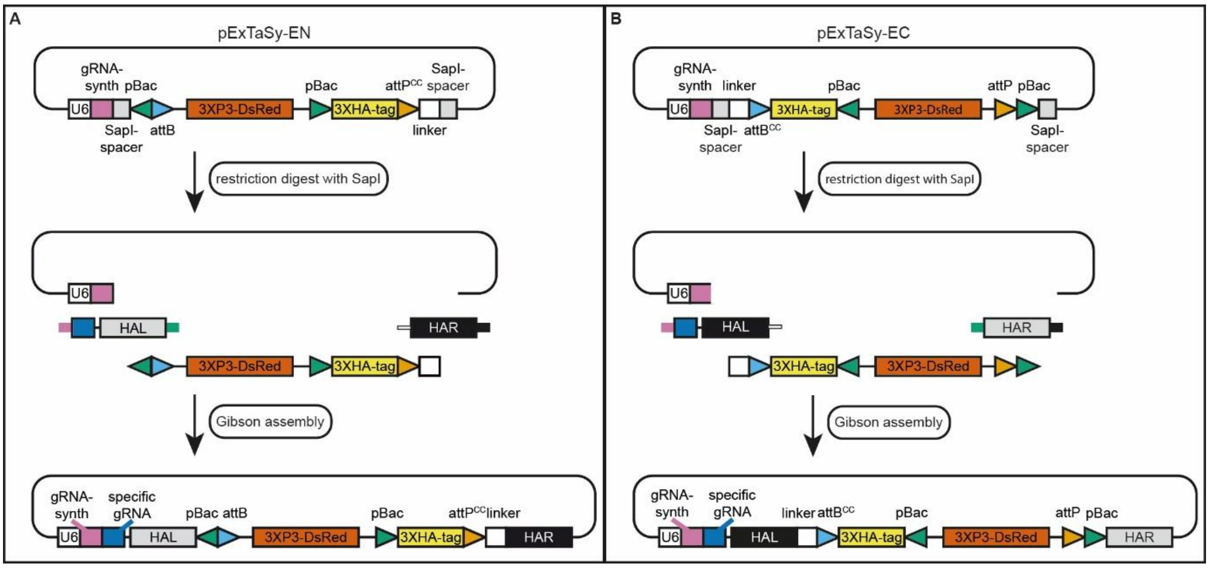

**Figure S1: Cloning procedure for ExTaSy transgenesis plasmids.** Restriction digest with SapI removes two spacer sequences, generating two fragments containing the vector backbone and the transgenesis cassette, respectively. A four-fragment Gibson assembly with two synthetic DNA fragments containing the left (HAL) and right (HAR) homology arms is used for cloning of the final plasmid. Due to different order of elements in the transgenesis cassette between *pExTaSy-EN* (A) and *pExTaSy-EC* (B), different overhangs for Gibson assembly are required (color-coded). For description of elements see Fig. 1.

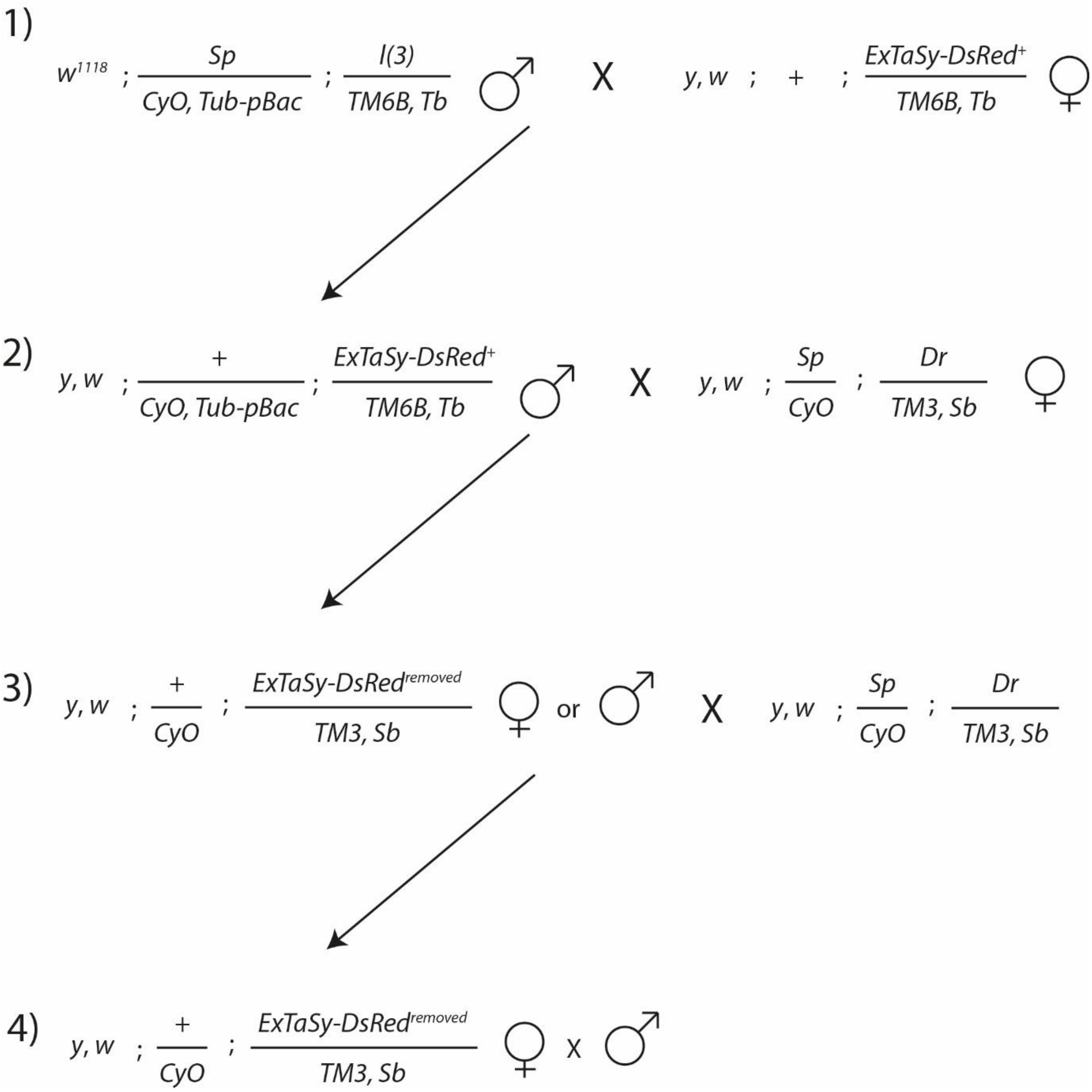

**Figure S2: Crossing scheme for transgenic marker removal.** The crossing scheme shows how the transgenesis marker can be removed from an ExTaSy insertion on *chr3* while keeping the genetic background the same. It works in an analogous fashion for insertions on other chromosomes but may require different balancer chromosomes. The Drosophila line expressing piggyBac transposase (pBac) is BDSC_8285 (see Table S1). This line is crossed to the ExTaSy line (1) and balanced male offspring with *CyO* marker are selected. These are crossed to a balancer line with second and third chromosome balancers (2). Offspring are then screened for loss of the DsRed transgenic marker and individually crossed to a double balancer (3). Balanced offspring can then be used to establish a breeding stock (4). PCR of the gene locus and Sanger sequencing should be used to confirm correct marker removal.

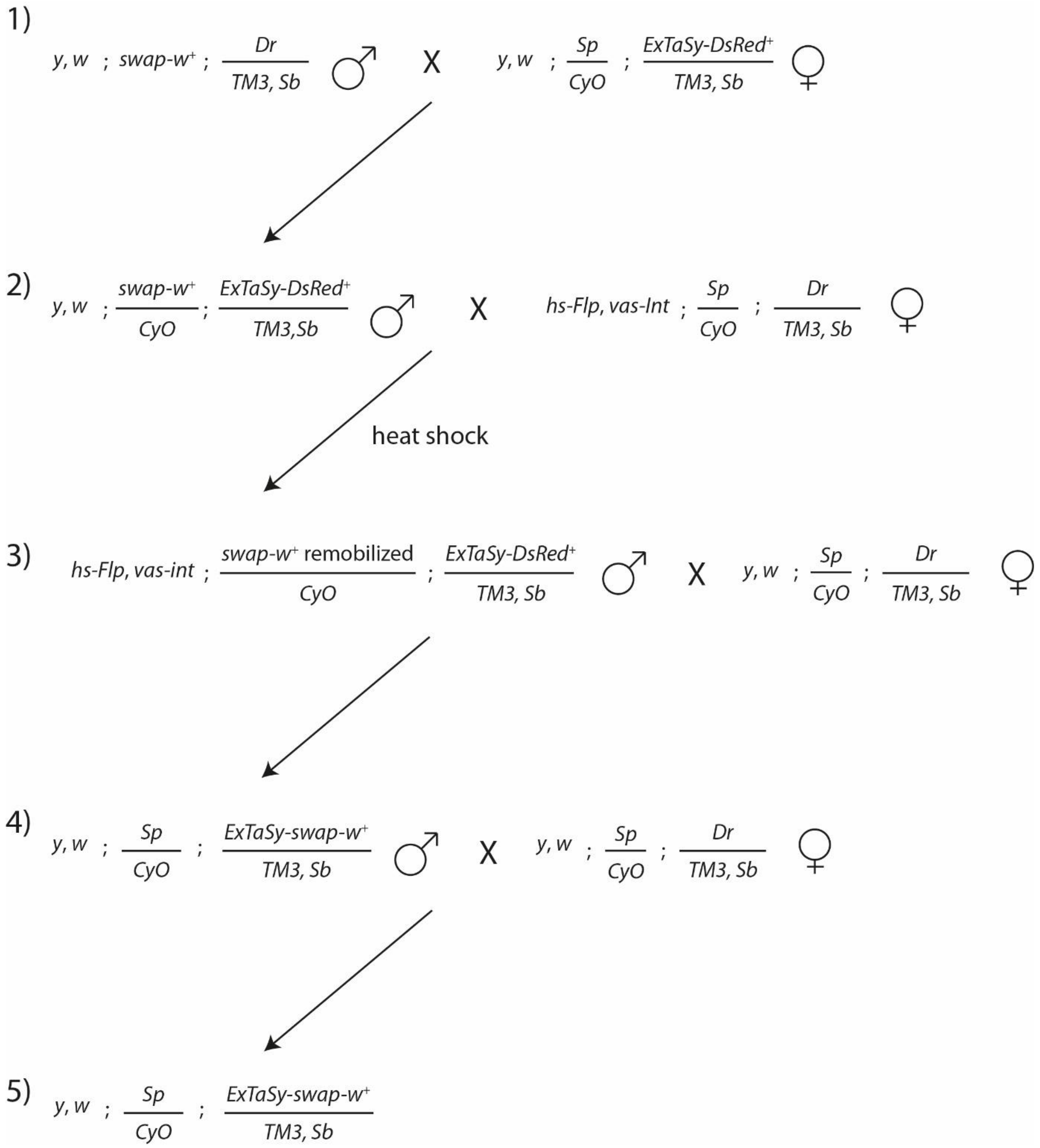

**Figure S3: Crossing scheme for tag exchange via RMCE.** The crossing scheme shows how the 3XHA tag can be exchanged for another tag at an ExTaSy insertion on *chr3* while keeping the genetic background the same. It works in an analogous fashion for insertions on other chromosomes but may require different balancer chromosomes. First, the primary tagged construct and the swap construct are brought together through a fly cross (1), then the enzymes for swap construct remobilization (Flippase) and RMCE (ΦC31 integrase) are added in a second cross (2). Offspring are heat-shocked within 48 h of egg lay for 1 h at 37 °C. The heat shock may be repeated after 24 and 48 h. Developing male offspring are then screened for mosaic expression of w in the eyes, which indicates successful remobilization of the swap construct. They are crossed to a double balancer line (3) and offspring are screened for loss of DsRed expression and gain of w expression (i.e., red eyes). They are again crossed to a balancer line (4) to create a breeding stock (5). PCR of the gene locus and Sanger sequencing should be used to confirm the tag swap.

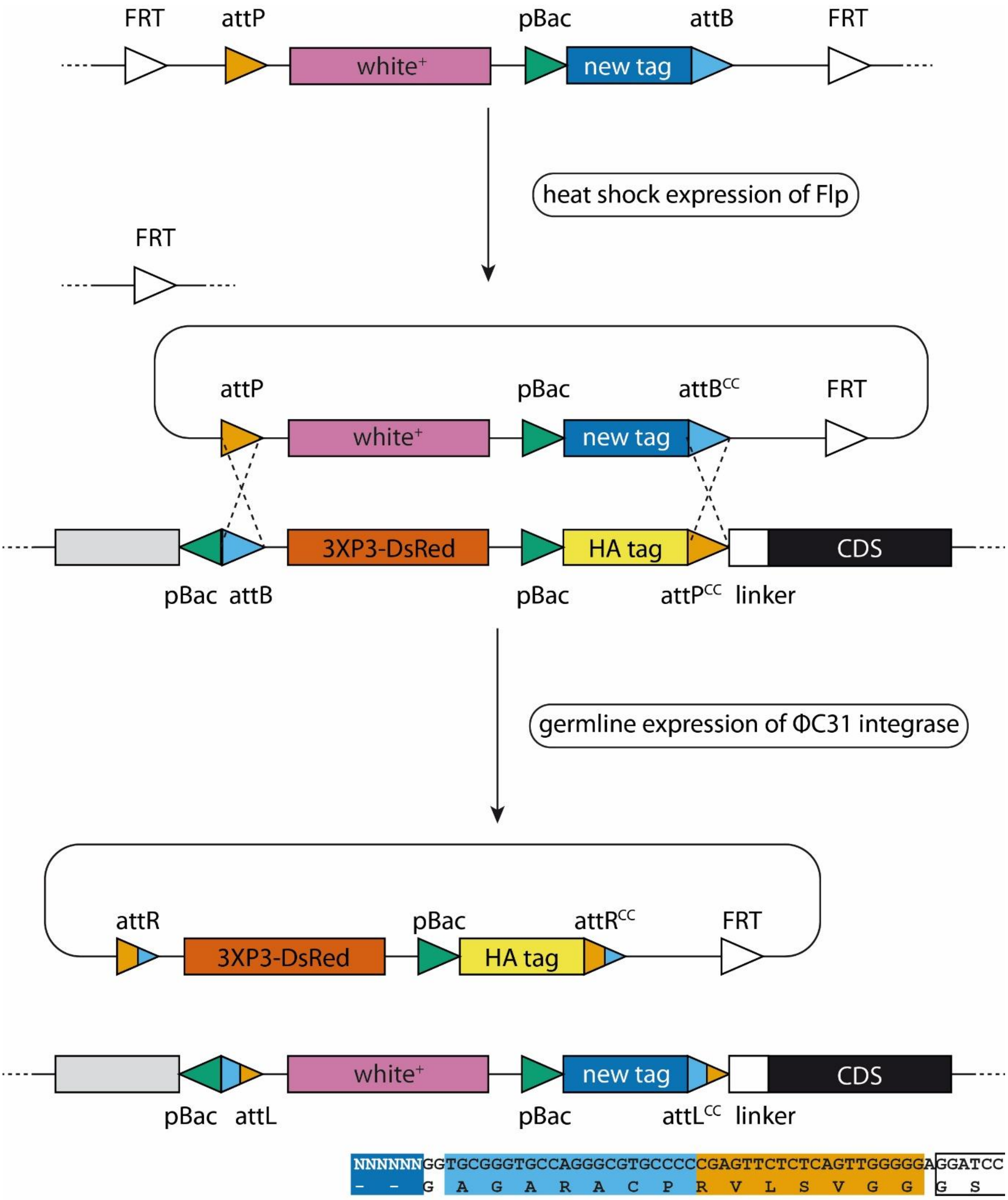

**Figure S4: Tag exchange for an N-terminally tagged gene.** Schematic of tag exchange for the N-terminus. The SwapSy construct is flanked by FRT sites. Flippase induces remobilization, leaving a single FRT site in the genome. ΦC31 integrase catalyzes recombination between the *attP*/*attB* and *attB*$^{CC}$/*attP*$^{CC}$ sites in the SwapSy and ExTaSy constructs. The exchange of tags and markers is unidirectional and leads to the formation of *attL* sites. The *attL*$^{CC}$ site links the new tag to the linker (nucleotide and amino acid sequences indicated below).

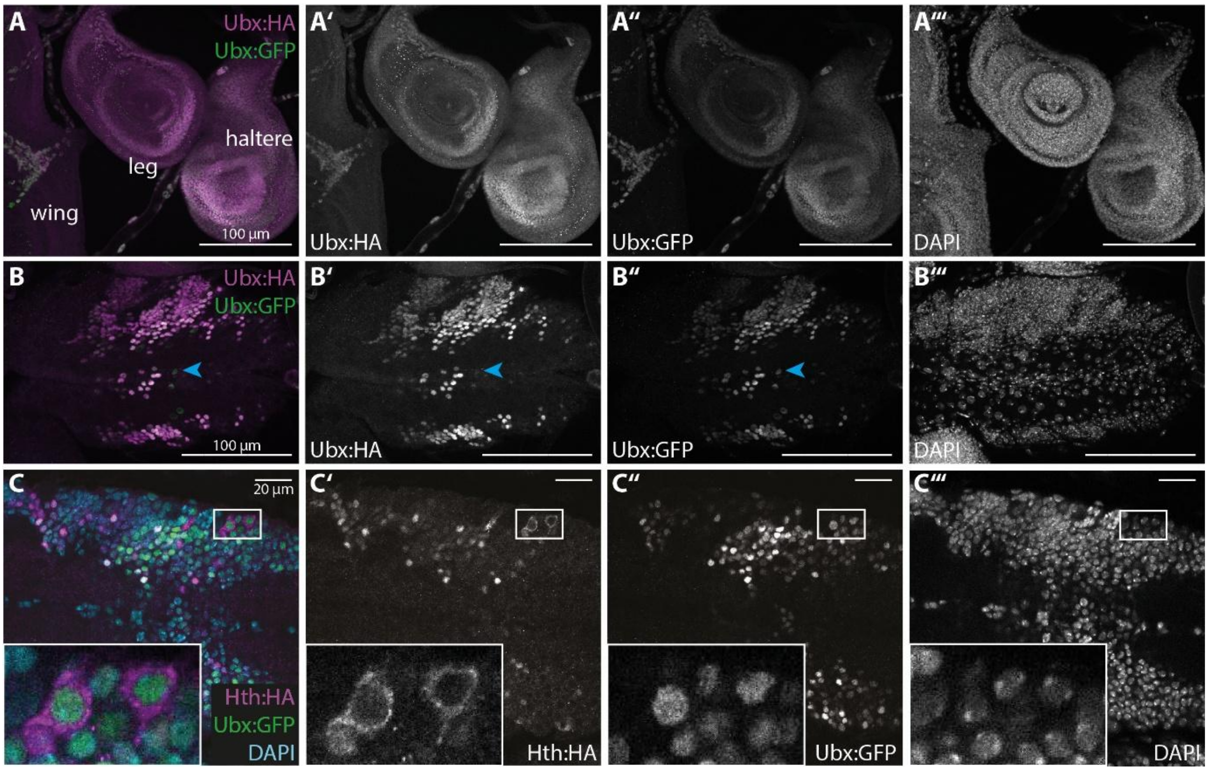

**Figure S5: Co-immunostaining of 3XHA- and sfGFP-tagged ~~Ubx~~ proteins.** Offspring of a cross between fly stocks with a C-terminally sfGFP-tagged *Ubx* allele (after tag exchange) and an N-terminally 3XHA-tagged allele (after marker removal). (A-A''') In haltere and third leg imaginal discs, there appears to be perfect overlap between the two stainings. (B-B''') Most cells in a ventral nerve chord in offspring of the same cross as in (A) show overlapping expression. Few cells only show staining for the sfGFP tag (arrowhead). Note that only a subsection of the ventral nerve chord is shown for better visibility. Anterior is to the left. (C-C''') Immunostaining of HA-tagged Hth (magenta) and sfGFP-tagged Ubx (green) in the ventral nerve chord. Anterior is to the left. DAPI staining in cyan shows nuclei. Cells in the periphery express both Ubx and Hth, but while Ubx is nuclear localized, Hth remains cytoplasmic (magnified in the inlay).
